# Structural Dynamics of the Dopamine D_2_ Receptor with a Non-Basic Ligand

**DOI:** 10.64898/2026.02.09.704831

**Authors:** Qinxin Sun, Guodong He, Damian Bartuzi, Andrea G. Silva, Ewa Kędzierska, Piotr Stępnicki, Anna Adamus, Katarzyna M. Targowska-Duda, Tomasz M. Wróbel, Marián Castro, Jens Carlsson, Xiangyu Liu, Agnieszka A. Kaczor

## Abstract

Progress in understanding protein-ligand interactions is revolutionizing drug design, especially for G protein-coupled receptors (GPCRs), which are targets for 35% of marketed drugs.^1^ The dopamine D_2_ receptor (D_2_R) represents a key drug target in schizophrenia and Parkinson’s disease.^2^ While structural studies have clarified its interactions with classical ligands, the behavior of atypical, non-basic ligands like D2AAK2 remains unclear. Notably, D2AAK2 shows strong selectivity for D_2_R over the closely related D_3_R, despite identical binding pocket composition. Here, we present a cryo-EM structure of D2AAK2 bound to D_2_R, showing that aspartate 3.32 serves as the main anchoring point, even though the compound lacks a basic nitrogen atom. Using enhanced sampling molecular dynamics simulations and experimental approaches, we uncover a complex binding energy landscape. Simulations suggest that D2AAK2 receptor subtype selectivity between identical binding sites arises from different energy barriers for their conformational changes. Non-basic ligands offer advantages such as better brain penetration and improved pharmacokinetics.^3,4^ This study provides the first structural insights into a non-basic ligand targeting D_2_R, paving the way for developing more effective, selective drugs.

## Main

G protein-coupled receptors (GPCRs) are one of the largest and most diverse families of membrane proteins, playing a central role in mediating physiological responses to a myriad of extracellular stimuli, including neurotransmitters, hormones, odorants, or photons.^5,6^ Their broad tissue distribution and involvement in numerous signaling pathways render GPCRs crucial targets for therapeutic intervention, with approximately one third of all FDA-approved drugs acting through these receptors.^7^ A key determinant of interactions between aminergic GPCRs and their orthosteric binders is the net charge of the ligands, due to Asp^3.32^ (Ballesteros-Weinstein numbering^8^) being a common feature of their orthosteric binding pockets. Virtually every known aminergic GPCR-targeting drug contains a protonatable nitrogen atom, which is positively charged under physiological conditions and capable of electrostatically interacting with the conserved aspartate. However, there is increasing evidence that neutral molecules can bind just as well, despite the absence of positively charged moieties in their structures.

Non-basic, or neutral, GPCR ligands have several key advantages over classical positively charged compounds. They are characterized by favorable pharmacokinetic properties, including oral bioavailability, metabolic stability, and reduced susceptibility to efflux transporters (eg. P-glycoprotein).^3^ They also have a limited probability of interacting with hERG potassium channels and a decreased risk of cardiotoxicity.^3,4^ Most importantly, they often display significant selectivity, even between nearly identical receptor subtypes, which can limit off-target effects and minimize the side effects of potential drugs.^9^

Indeed, the selectivity between closely related D_2_ and D_3_ receptors has been a persistent challenge in dopaminergic drug discovery, owing to the high degree of sequence and structural similarity within their orthosteric binding sites.^10,11^ Recent studies have elucidated how subtle differences, particularly secondary binding pockets within the extracellular loop region, can be exploited by appropriately designed ligands to enhance selectivity.^12^ Computational and structure-based drug design approaches, such as molecular dynamics (MD) simulations and machine learning-guided quantitative structure-activity relationship (QSAR) modeling or mutagenesis studies, have further highlighted features that confer selectivity for one subtype over the other.^13,14^ These findings, however, focus on highly exploited molecules that contain a protonatable nitrogen atom.^15^ Charge-neutral ligands can overcome some of the selectivity barriers historically associated with basic dopaminergic compounds by engaging with unique conformational states or by interacting preferentially with non-conserved residues. Despite this, there is a gap in understanding how GPCRs accommodate these molecules due to the lack of sufficient structural data. A detailed understanding of GPCR structure is essential for deciphering ligand recognition and receptor activation mechanisms.

Here, we present the first detailed description of a non-basic ligand bound to the D_2_ receptor (D_2_R). Using cryo-electron microscopy and molecular modeling, we define the structural features underlying this interaction. Our results reveal a distinct binding mode for non-basic ligands of aminergic receptors, as well as the basis for selectivity towards the D_2_R. These findings provide a structural framework for the development of selective ligands with properties beyond those accessible to classical charged molecules.

### Pharmacological characterization of D2AAK2

Our group discovered D2AAK2 (Fig. 1a) through structure-based virtual screening targeting D_2_R ligands.^16^ The compound exhibited medium to high nanomolar affinity for human cloned D_2_R in radioligand binding assays.^16,17^ Notably, D2AAK2 lacks a protonatable nitrogen atom, which is typically required to form an ionic interaction with Asp^3.32^ - a hallmark feature of orthosteric ligands targeting aminergic GPCRs. This unusual structural profile raised the question of whether D2AAK2 binds in an orthosteric or allosteric manner.

**Fig. 1:**
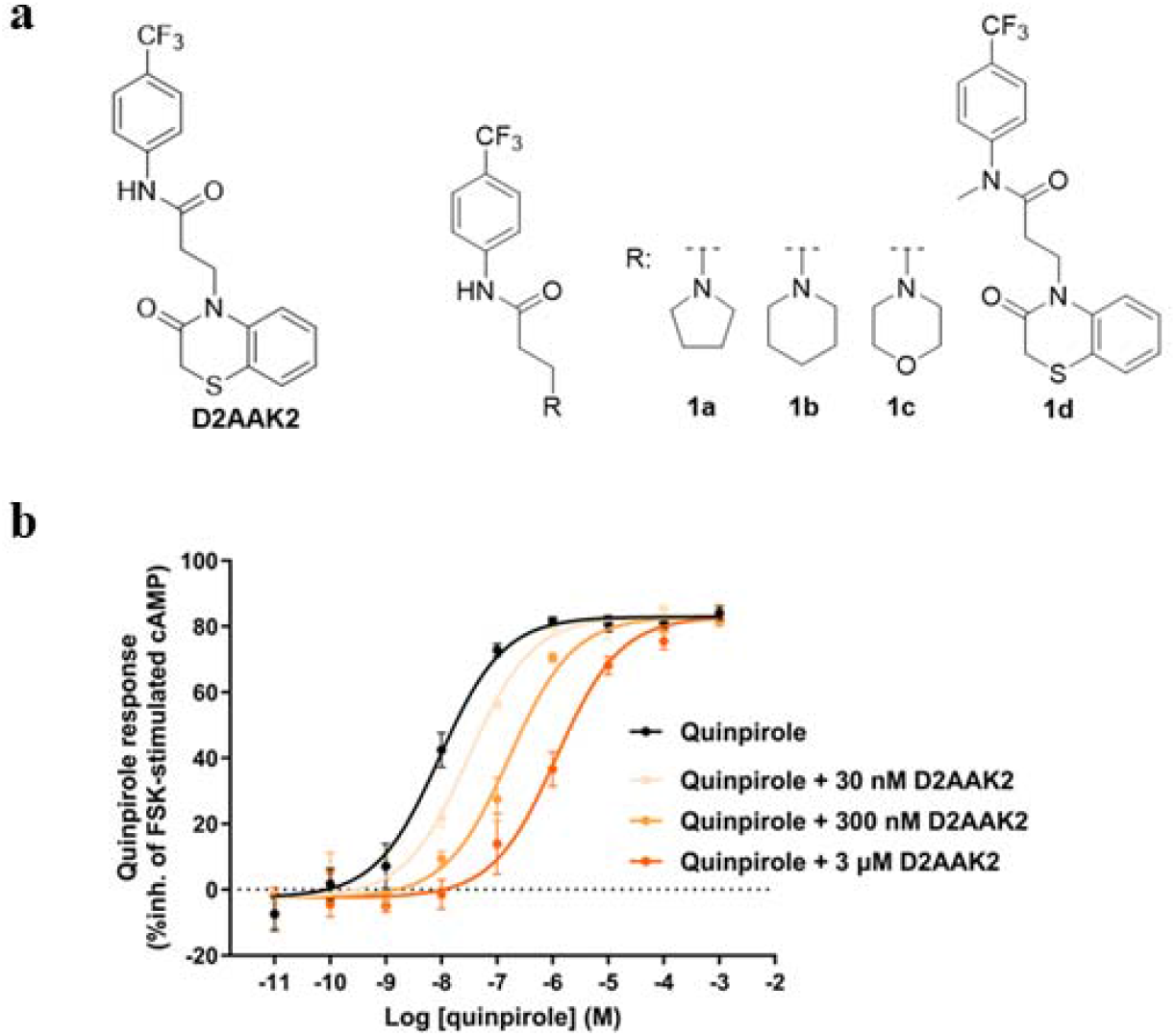
D2AAK2 is a highly selective, non-basic D_2_R antagonist. **a**, Chemical structure of D2AAK2 and its analogs. **b**, Concentration-response curves for quinpirole in the absence or presence of the indicated concentrations of D2AAK2, in cell-based functional assays of cAMP signaling at D_2_R. Data from individual experiments were expressed as % of inhibition (%inh.) of forskolin (FSK)-stimulated cAMP production and combined. Data points represent the mean ± SEM of three independent experiments performed in duplicate or triplicate. Curves drawn through the data points represent global fit to modified Gaddum/Schild EC_50_-shift model of competitive antagonism (Schild slope unconstrained, shared for all data sets) (fitting parameters provided in Supplementary Table S1).

The investigation of the selectivity profile of D2AAK2 at other aminergic receptors revealed its high selectivity for D_2_R over dopamine D_1_ and D_3_, serotonin 5-HT_1A_, 5-HT_2A_ and 5-HT_7_, histamine H_1_, and muscarinic M_1_ receptors (Table 1 and Supplementary Fig. S1a-h).^16,17^ Notably, the pronounced selectivity of D2AAK2 for D_2_ over D_3_ receptors – despite the nearly identical orthosteric binding pockets of these subtypes – suggests the involvement of an alternative, previously uncharacterized binding pocket. In functional assays of D_2_-mediated modulation of cAMP signaling, D2AAK2 antagonized dopamine response and elicited concentration-dependent dextral displacements of quinpirole concentration-response curves, a behavior compatible with competitive antagonism.^16^ However, negative allosteric modulators (NAMs), which decrease orthosteric ligand affinity without affecting its efficacy, would display a similar profile when the saturation of the allosteric binding site is not reached at the concentrations used in the assays.

**Table 1.** Affinity (expressed as equilibrium dissociation constant (*K*_i_) or % of inhibition of radioligand specific binding (%inh.)) of D2AAK2 at the indicated human cloned receptors.

| $K_i$ (nM) or % inh. at 10 μM | | | | | | | |
| --- | --- | --- | --- | --- | --- | --- | --- |
| *hD*<sub>2</sub> | *hD*<sub>1</sub> | *hD*<sub>3</sub> | *h5-HT*<sub>1A</sub> | *h5-HT*<sub>2A</sub> | *h5-HT*<sub>7</sub> | *hH*<sub>1</sub> | *hM*<sub>1</sub> |
| 58-321<sup>a</sup> | 2%<sup>b</sup> | 0%<sup>b</sup> | 13480<sup>b</sup> | 6%<sup>b</sup> | 26% | 5% | 2773 |

<sup>a</sup> range of $K_i$ values obtained for D2AAK2 samples obtained from different sources or preparation methods.<sup>16,17</sup>
<sup>b</sup> data from<sup>16</sup>.

We analyzed how D2AAK2 modulates quinpirole responses in cAMP assays at D_2_R. Quinpirole concentration-response curves, measured in the absence or presence of three concentrations of D2AAK2 (n = 3), were globally fitted to a four-parameter logistic model with shared maximum and slope, as required for Schild analysis. Allowing these two parameters to vary across curves did not improve the fit (extra sum-of-squares F-test (α = 0.05): F(6, 91) = 0.868, p = 0.522). Consistently, Akaike’s Information Criterion corrected for small sample size (AICc) favored the shared-parameter fitting (ΔAICc = 17.5). Based on this result, we fitted data globally to modified Gaddum/Schild EC_50_-shift model^18^ that would accommodate such behavior^19,20^ (preferred model over four-parameter logistic model (variable slope); ΔAICc = 2.336) (Fig. 1b). This retrieved a Schild slope value of 0.867 with 95% CI 0.728–1.02 that includes the standard value of 1.0. Hence, we could conclude that the Schild slope would not differ substantially from unity, consistent with a competitive antagonism mechanism. In fact, setting the Schild slope as a shared, unconstrained parameter did not fit the data significantly better than constraining it to unity (F-test (α = 0.05): F(1, 101) = 3.030, p = 0.0848). Therefore, we re-fitted the data by constraining the Schild slope to 1.0, which allowed us to extract p*K*_b_ value of D2AAK2 for D_2_R obtained from our functional assays as equal to p*A*_2_. *K*_b_ value was 21.06 nM (95% CI 3.02 nM–34.41 nM; n = 3), very close to the value previously reported for D2AAK2 obtained by linear Schild regression analysis.^16^

In line with a competitive antagonism mechanism − and contrary to what would be expected for a NAM − D2AAK2 (10 µM) did not alter the dissociation kinetics of the orthosteric radioligand [^3^H]-spiperone in a single experiment. Fitting of the data to one phase exponential decay model, allowing the dissociation rate constant to vary between data sets, did not improve the fit compared with a global fit with this parameter shared (F-test (α = 0.05): F(1, 25) = 0.005, p = 0.944) (Supplementary Fig. S2). Hence, global fit retrieved a dissociation rate constant for [^3^H]-spiperone from D_2_R of *k*_off_ = 0.0228 min^-1^ (95% CI 0.0197– 0.0262; n = 1), in good agreement with values previously reported for unlabeled spiperone in D_2L_R cell membrane preparations from competition binding kinetic assays (0.038 min^-1^).^21^ Nevertheless, because allosteric modulation can be probe-dependent, the choice of radioligand may limit our ability to detect certain NAM effects. In conclusion, our *in vitro* assays are consistent with D2AAK2 behaving as a highly selective, competitive D_2_R antagonist at the concentrations used.

### Cryo-EM structure of D2AAK2-D_2_R complex

To gain a deeper understanding of the molecular mechanism through which D2AAK2 interacts with D_2_R, we employed cryo-EM to determine the structures of the inactive state D_2_R bound to D2AAK2. We utilized an engineered human D_2_R construct with thermostabilizing mutations (I122^3.40^A, L375^6.37^A, and L379^6.41^A) and T4L insertion into the third intracellular loop (ICL3),^22^ along with a previously reported Fab fragment, Fab3089,^23^ for structure determination. We successfully solved the structure of D_2_R-D2AAK2 complex, with an overall resolution of approximately 2.99 Å (Fig. 2a and Supplementary Fig. S3). The clear electron density map allowed unambiguous modelling of the ligand and most parts of the receptor and the Fab, with the exception of ICL2 (141-147), the T4L fused to ICL3, and the constant domain of Fab3089 due to conformational flexibility.

**Fig. 2:**
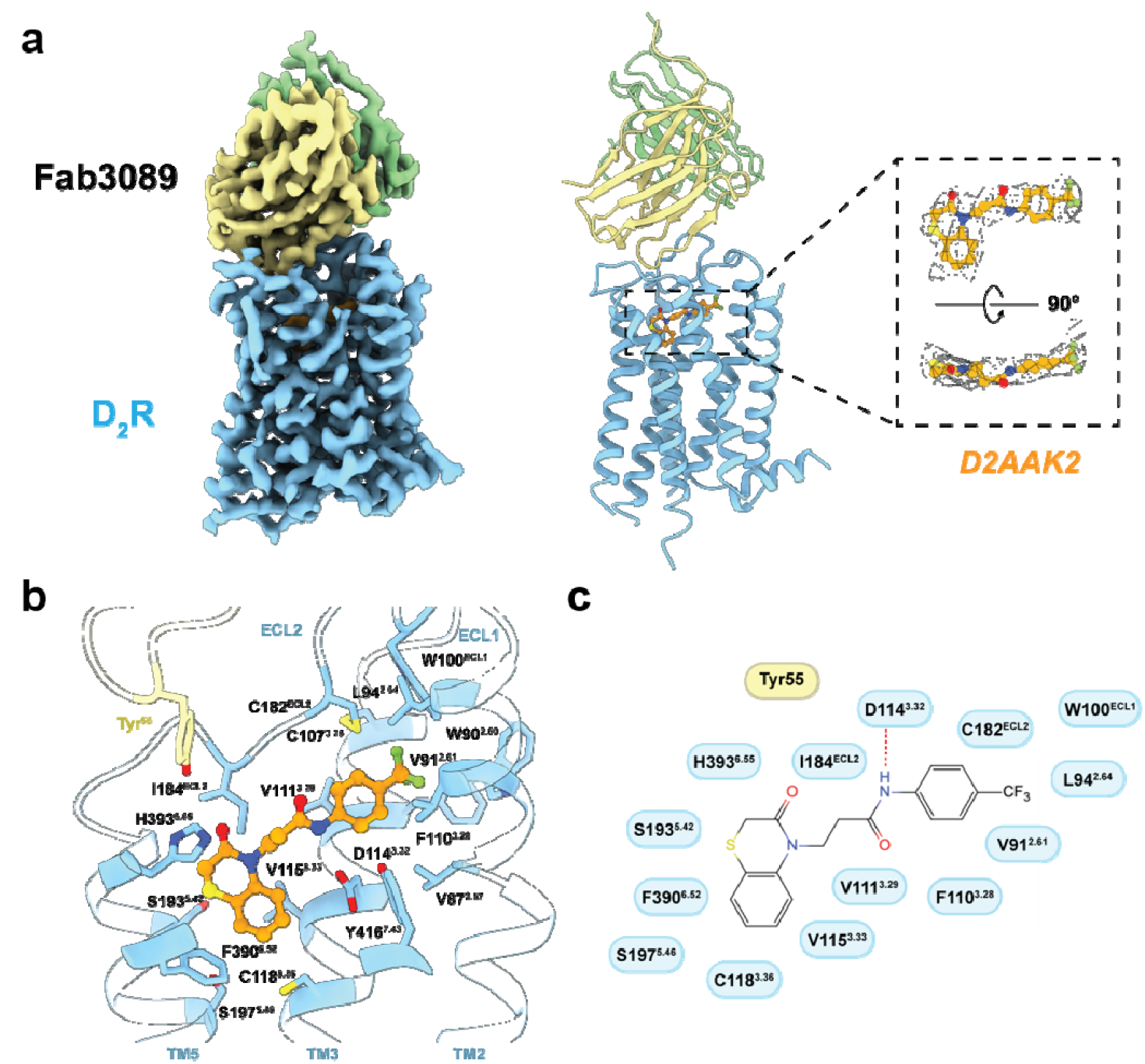
Cryo-EM structure of D2AAK2-bound D 2 R-Fab3089 complex. **a**, Overall structures of D 2 R bound with D2AAK2. From left to right: electron density map, coordinate model, and a zoom-in view of the electron densities for the ligand. **b**, Interactions between D2AAK2 and D_2_R. **c**, Diagram of the interactions between D_2_R and D2AAK2.

The complex structure provided a clear snapshot of how D2AAK2 binds to the orthosteric pocket of the receptor. In contrast to other D_2_R inverse agonists such as risperidone, haloperidol, and spiperone, which deeply penetrate and occupy the bottom hydrophobic cleft of the binding pocket, D2AAK2 adopts a more horizontal binding mode within a shallower region (Supplementary Fig. S4). The ligand forms extensive interactions with residues from TM2, TM3, TM5, TM6, TM7, ECL1 and ECL2. The benzothiazinone fragment of D2AAK2 fits into a pocket near TM5, while the trifluoromethylphenyl moiety extends toward the TM2 side, as will be discussed in more detail in subsequent sections.

D2AAK2 lacks a protonatable nitrogen atom - a key element of the classical D_2_R ligand pharmacophore (Fig. 1a). In canonical dopamine receptor ligands, the protonated nitrogen typically forms a salt bridge with Asp^3.32^, a strictly conserved residue in aminergic receptors (Fig. 3a-c). In the case of D2AAK2, the amide group instead forms a hydrogen bond with this conserved aspartate residue (Fig. 2b, c). This hydrogen bond is critical for ligand binding, as a close analog of D2AAK2 - differing only by methylation of the amide group (compound **1d**) shows no detectable affinity for D_2_R (Fig. 1a; Supplementary Table S2). Derivatives with the lactam ring replaced with cyclic amines further confirm that the moiety is not positioned favorably to form salt bridge with the conserved aspartate (compounds **1a-1c**) (Fig. 1a; Supplementary Table S2).

**Fig. 3:**
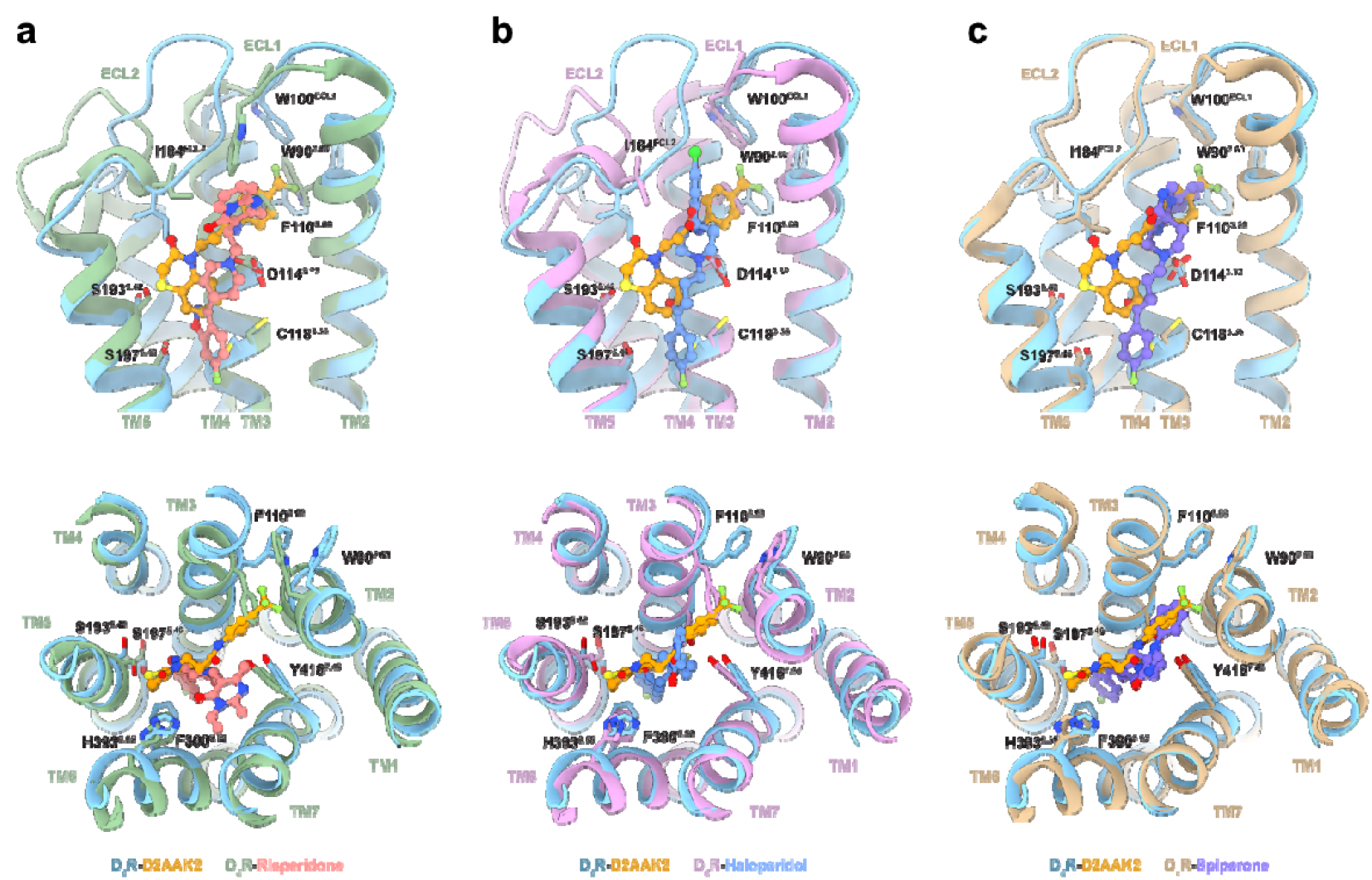
Comparison of D_2_R bound with D2AAK2 a) risperidone, b) haloperidol, c) spiperone. Compared to the D_2_R complexes bound with agonist bromocriptine or dopamine, the D2AAK2-bound D_2_R adopts a canonical inactive conformation, characterized by well-established structural features of GPCR inactivation. Key residues and motifs involved in receptor activation remain in their inactive configurations: Trp386^6.48^ retains its upward position (Supplementary Fig. S5c), the PIF motif stays compact (Supplementary Fig. S5d), and TM6 adopts an inward conformation (Supplementary Fig. S5a, b). While ECL2 is in proximity to the top of the ligand binding pocket in both active and D2AAK2-bound inactive states, the EBP of D2AAK2 is also disrupted in the agonist bound structures, due to the outward rotation of Trp100^ECL1^ and inward displacement of Phe110^3.28^ (Supplementary Fig. S5e, f).

Though D2AAK2 is not deeply inserted into the bottom hydrophobic clefts of the ligand binding pocket (Supplementary Fig. S4) unlike classical D_2_R antagonists, it still forms extensive interactions with the receptor (Fig. 2b,c). The benzothiazinone fragment occupies a subpocket formed by residues from TM5 and TM6, including Val115^3.33^, Ser193^5.42^, Ser197^5.46^ and Phe390^6.52^. Notably, in comparison with risperidone-, haloperidol-, and spiperone-bound D_2_R structures, the side chains of Ser197^5.46^ and Cys118^3.36^, which are located beneath the benzothiazinone fragment, flip toward the ligand in the D2AAK2-bound structure. This conformational change likely results from D2AAK2’s more superficial binding mode. Furthermore, unlike the aforementioned inverse agonists, which engage in hydrophobic interactions that stabilize the downward displacement of the toggle switch residue Trp386^6.48^, D2AAK2 lacks direct interaction with this key residue and does not induce the characteristic downward movement of Trp386^6.48^.

The trifluoromethylphenyl moiety of D2AAK2 occupies an extended binding pocket (EBP) within the cleft formed by TM2, TM3, ECL1, and ECL2. This EBP is previously identified in the D_2_R-spiperone complex (Fig. 3c) but is absent in structures bound to risperidone or haloperidol (Fig. 3a, b). Key residues contributing to the EBP include Trp90^2.60^, Val91^2.61^, Leu94^2.64^, Trp100^ECL1^, Phe110^3.28^, Val111^3.29^, and Cys182^ECL2^, which form hydrophobic interactions with D2AAK2’s trifluoromethylphenyl group. Notably, the side chain of Phe110^3.28^ undergoes a conformational shift toward the membrane side to accommodate D2AAK2 and forms a π-stacking interaction with Trp90^2.60^ to stabilize the EBP architecture. In contrast, the EBP is occluded in risperidone- and haloperidol-bound D_2_R structures due to the inward rotation of Phe110^3.28^. Specifically, in the risperidone-bound structure, the inward orientation of Trp90^2.60^ and Trp100^ECL1^ further compresses the EBP space.

Additionally, similar to the spiperone-bound state, ECL2 covers the top of ligand-binding pocket, facilitating direct contacts between Ile184 and Cys182 on ECL2 and the ligand. In contrast, ECL2 extends away from the receptor core in both D_2_R-risperidone and D_2_R-haloperidol complexes. However, it is important to note that the ECL2 conformation might be influenced by the binding of Fab3089. Notably, Tyr55 of Fab3089’s heavy chain also inserts into the extracellular vestibule of the receptor and interacts with D2AAK2.

### Molecular basis of D2AAK2 binding and selectivity

To gain further understanding of D2AAK2 binding to D_2_R, we performed biased and unbiased MD simulations based on the cryo-EM structure. These calculations aimed at validating the binding mode of the ligand, characterizing interactions with binding site residues and solvent molecules, and understanding the unusual D_2_/D_3_ selectivity profile. In the first step, we evaluated the impact of the protonation state of Asp^3.32^ on D2AAK2 binding. Whereas dopamine and synthetic ligands of D_2_R are typically cations and form a salt bridge to Asp^3.32^, D2AAK2 is neutral and could potentially affect the protonation state of the side chain carboxylate. Although pK_a_ calculations with PropKa^24,25^ indicate prevalence of the charged form of the Asp^3.32^ residue, the calculated pK_a_ value is shifted from 5.8 in the apo receptor to 6.1 upon insertion of D2AAK2 in the binding site. The predicted value is thus close to 7, at which populations of both forms would be equally represented. For comparison, pKa values calculated for the 7DFP structure in an apo and spiperone-bound state drop upon this classical ligand binding from 5.4 to 3.0. However, unbiased MD simulations indicated high stability of the complex with a charged Asp^3.32^ side chain. In contrast, the interaction between D2AAK2 and the side chain carboxylate of Asp^3.32^ was disrupted in simulations of the neutral form. Thus, both MD simulations and PROPKA predictions support that charged Asp^3.32^ is prevalent in D2AAK2-bound D_2_R Analysis of sodium ion occupancies in the apo and holo systems using unbiased MD simulations revealed a slight increase in sodium ion occupancy at Asp^3.32^ in the ligand-bound receptor (Supplementary Fig. S6), which may further stabilize the complex.

The cryo-EM structure also revealed that the hydroxyl group of Tyr^55^ of the stabilizing antibody protrudes toward the orthosteric pocket, at a distance of 4 Å from D2AAK2. To assess if the stabilizing antibody affected the ligand binding mode observed in the cryo-EM structure, we performed funnel metadynamics (FM) simulations of the binding process. The ligand pose corresponding to the deepest free energy well nearly perfectly overlaps with the pose consistent with the cryo-EM data (Fig. 4b). In addition, the calculated binding free energy (−10.6 kcal/mol) agrees with the values calculated from the radioligand binding assay (between −10.3 and −9.0 kcal/mol). Therefore, FM shows that the lowest energy conformation of the receptor-D2AAK2 complex with no antibody bound agrees with the pose observed in the cryo-EM structure, indicating that the antibody did not affect the D2AAK2 binding. Water occupancy analysis in unbiased MD simulations showed that the volume occupied by the hydroxyl group of Tyr^55^ was replaced by a stable water molecule in simulations (Supplementary Fig. S6). The FM simulations also provided detailed insights into the binding process and roles played by particular ligand moieties. The binding free energy landscape unveils a broad valley, stretching from the initial ligand capturing point in an extracellular vestibule to the orthosteric site buried in the receptor interior (Fig. 4a). The path to the global free energy minimum leads through several metastable intermediate states (Fig. 4c). Notably, the first binding events involve the aromatic ring of the benzothiazinone moiety, that would not be possible in case of the inactive **1a, 1b** and **1c** derivatives as they do not contain a corresponding aromatic moiety, suggesting a potential importance of the ligand binding path for antagonist binding. These initial events involve interactions with cleft at ECL3, in agreement with the earlier findings by Dror et al. for the β_2_ adrenergic receptor.^26^

**Fig. 4:**
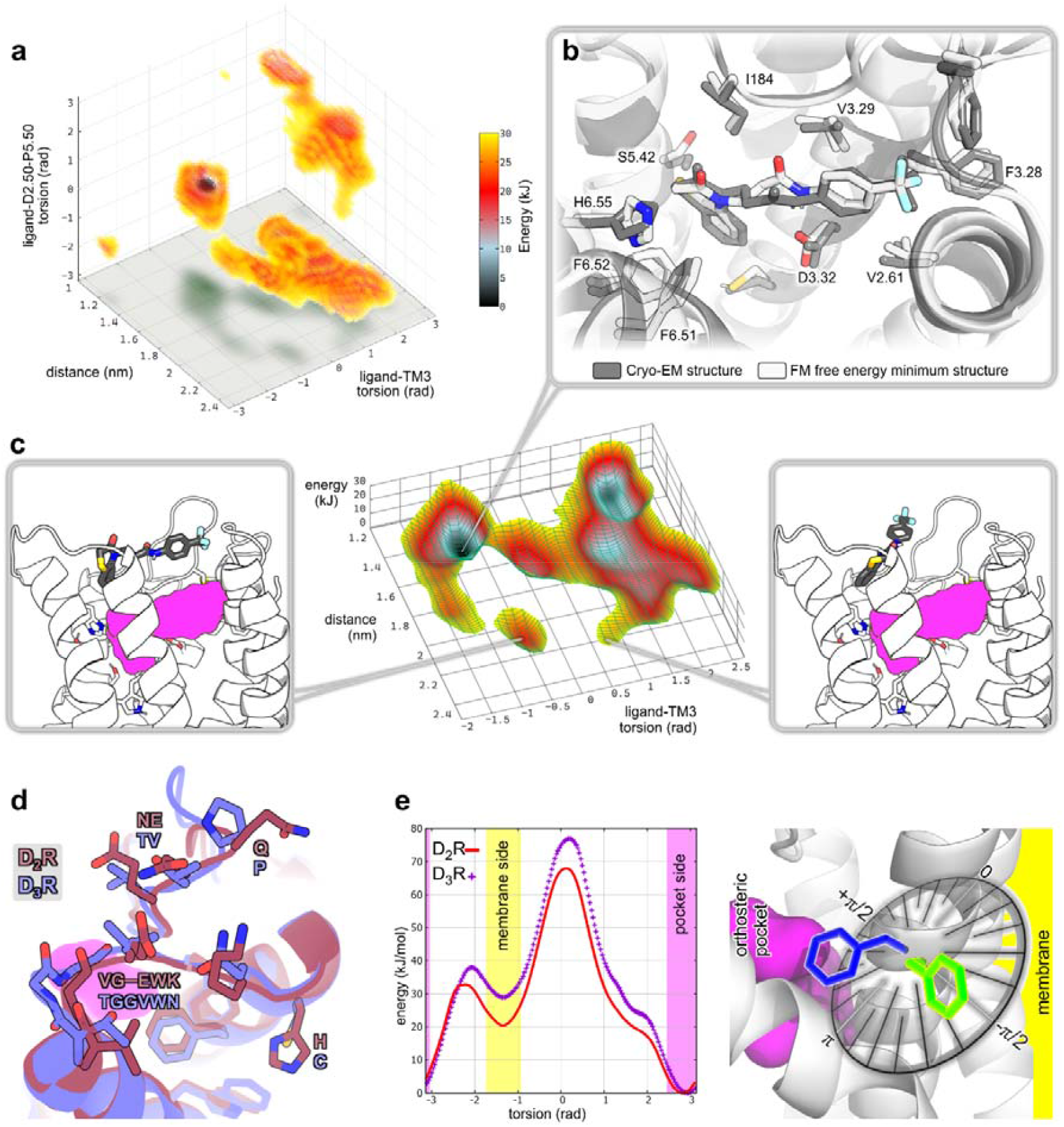
FM simulations of D2AAK2 binding to the D_2_R. **a**, The free energy landscape in three dimensions, presenting values of all three collective variables used in the funnel metadynamics simulations. **b**, D2AAK2 binding mode in the cryo-EM structure (gray) compared to the free energy minimum complex (white). **c**, D2AAK2-receptor initial binding events, pinned to their corresponding areas of FES. The orthosteric binding site is marked in purple in the structures. **d**, Comparison of D_2_R and D_3_R structures in the ECL1 and ECL2 region. The orthosteric binding site is marked in purple. **e**, Free energy profiles of Phe^3.28^ rotation in the D_2_R and D_3_R structures calculated using well-tempered metadynamics simulations.

The binding mode revealed by the cryo-EM structure does not explain the structural basis of D2AAK2 D_2_/D_3_ selectivity, as the residues of the orthosteric pocket directly interacting with the ligand are identical in both receptor subtypes. Since no difference in the binding pocket composition was identified, we turned our attention to the immediate surroundings. Analysis of experimental D_2_R structures showed that binding site residue Phe^3.28^ can occupy two main rotamers, and binding of D2AAK2 is only compatible with the outward rotamer, in which the side chain faces the membrane. Inspection of the Phe^3.28^ surroundings reveals several sequence variations between D_2_R and D_3_R, including a difference in ECL1 length, polarity of side chains, and presence or absence of flexible or rigid residues (Fig. 4d). These differences may indirectly influence the binding pockets in both receptor subtypes and affect the ability of Phe^3.28^ to switch the rotameric state. This conformational change may, in turn, affect the affinity of D2AAK2 for the different receptor subtypes. To assess the impact of the surrounding residues on Phe^3.28^ rotation, we performed well-tempered metadynamics with a single CV representing the phenylalanine’s χ_1_ torsion for both D_2_ and D_3_ receptor structures. The free energy profiles reveal differences in the Phe^3.28^ side chain behavior in the two receptors. In particular, there was nearly a 10 kJ/mol difference between the subtypes for the local minimum of the membrane-facing conformation of Phe^3.28^ and 5 kJ/mol for the rotamer-transition energy barrier height. These results suggest a higher likelihood of the binding site conformation enabling D2AAK2 binding in the D_2_ receptor (Fig. 4e), explaining the higher affinity for this receptor. Notably, while the outward Phe^3.28^ conformation in D_2_R was observed in the spiperone complex (PDB ID: 7DFP, Fig. 3c), none of the available experimental D_3_R structures include this feature.

### D2AAK2 effect on amphetamine-induced animal hyperactivity in male mice

Several doses of D2AAK2 (6.5, 12.5, 25, and 50 mg/kg, i.p.) were tested for their effect on mouse spontaneous locomotor activity. The results indicated that the 50 mg/kg dose significantly reduced locomotor activity, as shown in Supplementary Fig. S7. In the next step, three doses of the studied compound (6.5, 12.5, and 25 mg/kg) that had no effect on mouse locomotor activity were selected to assess their impact on amphetamine-induced hyperactivity. Statistical analysis using two-way ANOVA, presented in Supplementary Table S3, showed significant D2AAK2 pretreatment and amphetamine treatment effects, and ANOVA interaction between both treatments.

More specifically, the post hoc Tukey’s test confirmed a significant increase in the distance traveled by mice after amphetamine treatment compared to vehicle-treated controls (Fig. 5) and a significant decrease in distance traveled by animal after co-injection of D2AAK2 (12.5 and 25 mg/kg) with amphetamine when compared with amphetamine treatment alone (Fig. 5). No significant effect was observed with the 6.5 mg/kg dose of the compound (Supplementary Table S3).

**Fig. 5:**
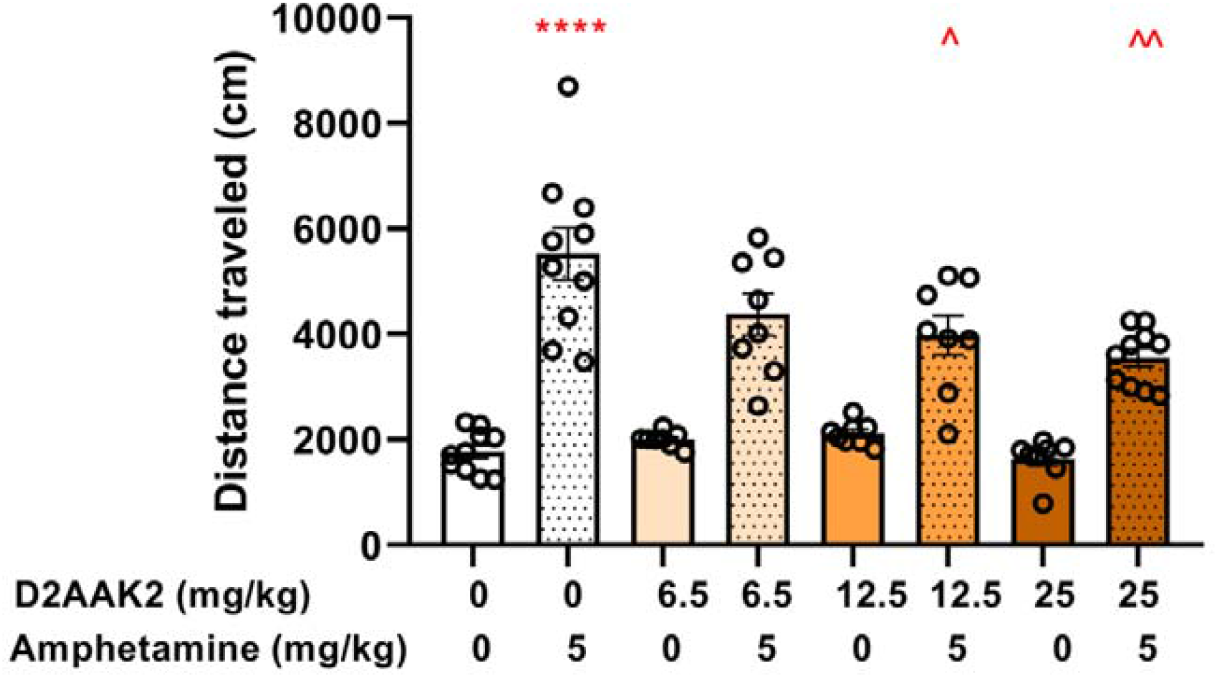
Influence of D2AAK2 (6.5, 12.5, and 25 mg/kg, i.p.) on amphetamine-induced hyperactivity determined in male mice. Mice in each group were injected with D2AAK2 (n = 8-10), amphetamine (n = 10), vehicle (n = 10), and D2AAK2 co-injected with amphetamine (n = 8-10). Mice were placed into the locomotor activity chamber 30 min after D2AAK2/vehicle treatment (i.e., immediately after amphetamine/vehicle injection). The values represent the mean ± SEM of the distance traveled by animals from each treatment group. The post hoc test revealed that there is an increase in distance traveled after amphetamine injection (**** p < 0.0001) and a decrease in distance traveled after co-injections of 12.5 or 25 mg/kg D2AAK2 with amphetamine when compared to amphetamine treated mice (^p = 0.0489 for 12.5 mg/kg D2AAK2 and ^^ p = 0.001125 for 25 mg/kg D2AAK2, two-way ANOVA followed by Tukey’s post hoc test).

## Discussion

The understanding of structural aspects of ligand-receptor interactions is crucial for drug design. In the case of aminergic GPCRs, this phenomenon is relatively well investigated for ionizable orthosteric ligands. In this context, the present study challenges the classical model of aminergic GPCR ligands binding^27^ by showing that a non-basic ligand, D2AAK2, can interact with D_2_R through a unique orthosteric mode. Our combined structural, computational, and pharmacological investigations provide a comprehensive picture of the binding dynamics and selectivity determinants for D2AAK2 at D_2_R. Although D2AAK2 does not have the protonatable nitrogen atom present in most aminergic ligands, our cryo-EM structure clearly shows that it occupies the orthosteric site of D_2_R and interacts with the conserved Asp3.32, which is the key anchoring point. This finding, supported by our molecular dynamics and funnel metadynamics simulations, show that non-basic ligands may display a nanomolar affinity thanks to their involvement in very specific interactions, partly mediated by hydrogen bonds and a network of water molecules and sodium ions.

Our results show that the microenvironment^28^ surrounding the receptor plays an essential role in stabilization of the binding pose of D2AAK2. Both water molecules and ions may have a structural role in proteins and take part in ligand binding and activation mechanisms of GPCRs.^29–32^. Apart from the well-established role of allosteric sodium located in the buried pocket at Asp^2.50^ residue,^33^ various studies indicate other possible roles played by these ions. In particular, recent work of Chan et al. suggests a ligand-stabilizing effect of sodium and magnesium ions in dopamine and opioid receptors, respectively.^34^ Interestingly, the authors suggest a possibility of sodium ion binding at His^6.55^, additionally coordinated by the quinolin-8-ol moiety of the MLS1547 ligand. As D2AAK2 displays zero net charge but is capable of binding to the charged binding site with nanomolar affinity, a possibility of a stabilizing role of a sodium ion was also investigated. Analysis of water and sodium ion occupancies allows us to conclude that sodium ions might stabilize the Asp^3.32^ conformation in the ligand-bound receptor. Furthermore, the conserved water molecules found in the cryo-EM map and verified by MD trajectories create a structural network that supports the proper orientation of both the ligand and key receptor residues.

Our molecular modeling studies make it possible to understand the dynamic process of ligand binding. Funnel metadynamics found a broad valley in the free energy landscape, characterized by a number of metastable states throughout the binding pathway. The most profound free energy basin aligns well with the binding pose seen through cryo-EM, but the landscape also shows temporary trapping in less ideal poses. A particularly intriguing finding is the distinct behavior of phenylalanine 3.28 (Phe^3.28^). Our data shows that rotation of its side chain is necessary to accommodate the trifluoromethylphenyl moiety of D2AAK2. Free energy calculations confirm that Phe^3.28^ rotates more easily in D_2_R in comparison to D_3_R, providing a possible explanation for the observed selectivity of D2AAK2 for D_2_R. In D_3_R subtle differences in the extracellular loops and neighboring residues (e.g., a Cys substitution for His^3.24^ and differences in the ECL1 and ECL2 sequence) may restrict such conformational flexibility and the energy barriers for phenylalanine reorientation are higher. These structural nuances may underlie the preferential binding of D2AAK2 to D_2_R, highlighting the potential of exploiting non-conserved dynamic features among receptor subtypes for selective drug design.

We also explored D2AAK2 analogs – compounds **1a** - **1d** – to further investigate the compounds’ binding mode. We introduced protonatable groups (compounds **1a**–**1c**) to enhance interactions with Asp3.32, but these changes led to a loss of D_2_R affinity. Methylation of the amide group (compound **1d**) also resulted in an inactive derivative which supported the importance of the amide hydrogen atom in the binding mode, proposed in our earlier modeling studies^16^ and confirmed by the Cryo-EM structure. These findings stress that even minor structural modifications can disrupt the balance of hydrophobic and aromatic interactions required for effective receptor binding, further validating the unique, binding paradigm of D2AAK2.

From a molecular pharmacology perspective, the functional assays suggest that D2AAK2 acts as a competitive, reversible antagonist at D_2_R. Furthermore, the *in vivo* behavioral studies, where D2AAK2 decreased amphetamine-induced hyperactivity, correlate with the receptor binding data and provide a functional link^35^ to our structural observations.

Non-basic D2AAK2 compound represents a promising hit for therapeutic development due to a number of benefits over classical ionizable ligands.^3^ Its neutral charge limits the risk of off-target interactions,^36^ but also improves pharmacokinetic properties, including metabolic stability and blood-brain barrier permeability. These are key features for drugs targeting central nervous system disorders such as schizophrenia and Parkinson’s disease, where precise receptor modulation is essential.^36^ Next, by avoiding the common drawbacks of basic ligands - such as nonspecific receptor activation and cardiotoxicity risks - D2AAK2 may have a safer drug profile and improved selectivity. Taken together, these properties emphasize the potential of D2AAK2 as a drug candidate for more effective and tailored therapies targeting D_2_R.

In summary, our work shows that non-basic ligands, in spite of lacking classical ionic interactions, can achieve high-affinity binding to aminergic receptors. It involves water-mediated hydrogen bonding and subtle interplay with ions, such as sodium ions. Our combined structural, computational, and pharmacological approach not only validates the orthosteric binding mode of D2AAK2 but also supplies the mechanistic insights into receptor subtype selectivity that could be useful for the design of future drugs with improved selectivity and pharmacokinetic profiles. Our model provides a novel scaffold upon which further research can be conducted towards selective GPCR ligands.

## Materials and methods

### D_2_R-T4L-D2AAK2 expression and purification

The expression and purification of the engineered D_2_R-T4L was performed following a previously reported method.^22^ Briefly, the receptor was expressed in Spodoptera frugiperda (Sf9) cells (Expression Systems) using the Bac-to-Bac Baculovirus Expression System (Invitrogen) for 48 hours. The collected cell pellets were lysed in the lysis buffer containing 20 mM HEPES pH=7.5, 10 mM MgCl_2_, 20 mM NaCl, 10 μM D2AAK2, and protease inhibitors. After centrifugation, the pellet was collected and solubilized in the solubilization buffer containing 20 mM HEPES pH=7.5, 800 mM NaCl, 1% (wt/vol) DDM, 0.1% (wt/vol) CHS, 10 μM D2AAK2, 20 μM imidazole pH=8.0, and protease inhibitors for 2 hours at 4 °C.

The supernatant of the solubilized cell membrane was obtained through centrifugation and was then incubated with pre-equilibrated Nickel resin for 2 hours. The resin was washed alternately with low salt buffer (20 mM HEPES, pH=7.5, 150 mM NaCl, 0.1% (wt/vol) DDM, 0.01% (wt/vol) CHS, 10 μM D2AAK2, and 20 μM imidazole, pH=8.0) and high salt buffer (20 mM HEPES, pH=7.5, 800 mM NaCl, 0.1% (wt/vol) DDM, 0.01% (wt/vol) CHS, 10 μM D2AAK2, and 20 μM imidazole, pH=8.0). The protein was eluted using Nickel Elution Buffer (20 mM HEPES, pH=7.5, 150 mM NaCl, 0.1% (wt/vol) DDM, 0.01% (wt/vol) CHS, 10 μM D2AAK2, 2 mM CaCl_2_, and 250 μM imidazole, pH=8.0), followed by loading into a pre-equilibrated M1 anti-FLAG column for further purification. The detergent was exchanged to 0.01% (wt/vol) LMNG and 0.002% (wt/vol) CHS on the column.

The receptor was eluted from the M1 anti-FLAG column using the M1 elution buffer (20 mM HEPES, pH=7.5, 150 mM NaCl, 0.01% (wt/vol) LMNG, 0.002% (wt/vol) CHS, 20 μM D2AAK2, 5 mM EDTA, and 0.2 mg/ml FLAG peptide), followed by size exclusion chromatography (SEC) with a final SEC buffer consisting of 20 mM HEPES, pH 7.5, 150 mM NaCl, 0.002% (wt/vol) LMNG, 0.0004% (wt/vol) CHS, and 20 μM D2AAK2. The purified protein was concentrated using a 50 kDa MWCO concentrator, flash-frozen in liquid nitrogen, and stored at −80°C for future use.

### Fab3089 expression and purification

The cDNA sequences of the Fab3089 heavy chain (Fab-H) and light chain (Fab-L) were subcloned into pcDNA3.1, with a signal peptide (MGWSCIILFLVATATGVHS) added to their N-terminus. Additionally, a 6×His tag was appended to the C-terminus of Fab-L for purification. For expression, plasmids of Fab-H and Fab-L were co-transfected into Expi293F cells using PEI. The cells were cultured in SMM 293-TI medium (Sino Biological Inc) at 37°C for 5 days after transfection.

After protein expression for 5 days, the cell culture medium was collected and loaded onto pre-equilibrated Nickel resin. The resin was washed with a buffer containing 20 mM HEPES (pH=7.5), 150 mM NaCl, and 20 mM imidazole (pH=8.0). Finally, Fab3089 was eluted from the resin using a buffer containing 20 mM HEPES (pH=7.5), 150 mM NaCl, and 250 mM imidazole (pH=8.0). Imidazole and other contaminants were further removed through size exclusion chromatography, with a final buffer consisting of 20 mM HEPES (pH 7.5) and 150 mM NaCl. The purified protein was concentrated using a 30 kDa MWCO concentrator, flash frozen in liquid nitrogen, and stored at −80°C for future use.

### D_2_R-T4L-D2AAK2-Fab3089 complex formation

To assemble the D_2_R-T4L-D2AAK2-Fab3089 complex, purified D2R-T4L-D2AAK2 was incubated with a 1.2 molar excess of Fab3089 for 1 hour at room temperature. Following incubation, the mixture was loaded onto a pre-equilibrated M1 anti-FLAG column to remove excess Fab3089. The complex was eluted from the resin using a buffer containing 20 mM HEPES, pH 7.5, 150 mM NaCl, 0.002% (wt/vol) LMNG, 0.0004% (wt/vol) CHS, 5 mM EDTA, 0.2 mg/ml FLAG peptide, and 20 μM D2AAK2. Subsequently, size exclusion chromatography was performed with a buffer consisting of 20 mM HEPES (pH 7.5), 150 mM NaCl, 0.002% (wt/vol) LMNG, 0.0004% (wt/vol) CHS, and 20 μM D2AAK2 for further purification. The purified protein was concentrated using a 50 kDa MWCO concentrator, flash frozen in liquid nitrogen, and stored at −80°C for cryo-EM sample preparation.

### Cryo-EM sample preparation and data collection

For the cryo sample preparation of D2R-T4L-D2AAK2-Fab3089 complexes, 4 µL of protein was loaded onto glow-discharged holey carbon grids (Quantifoil Au R1.2/1.3, 300 mesh). The grids were blotted for 4.5 seconds and flash-frozen in liquid ethane using a Vitrobot (Mark IV, Thermo Fisher Scientific). Cryo-EM data were collected on a Titan Krios G3i microscope equipped with a Gatan K3 Summit detector at the National Protein Science Research Facility Tsinghua Base in Beijing. Movie stacks were collected with a defocus range from −1.3 µm to −1.6 µm with a total dose of about 50 e-/Å^2^ using the AutoEMation program. Each stack was exposed for 2.56 seconds, and 32 frames were recorded per micrograph and aligned by MotionCor2^37^ and binned to a pixel size of 1.08 Å.

### Cryo-EM data process and structure determination (see Supplementary Fig. S3)

Dose-weighted micrographs were imported into cryoSPARC,^38^ and CTF parameters were estimated by patch-CTF. Then 1,812,899 particles were picked from 1384 micrographs by blob-picker. Two rounds of 2D classification were performed to reveal the clear feature of the GPCR-Fab complex, and particles showing all features of GPCR-Fab complex were selected to generate the initial model by ab-initio reconstruction. By multiple rounds of heterogeneous refinement, 563,944 particles were subjected and re-extracted. With heterogeneous refinement, non-uniform refinement and local refinement, the final cryo-EM map was obtained with a resolution of 2.99 Å, with the density of T4L missing because of its flexibility.

D2AAK2’s model and restraints were generated by eLBOW in Phenix1.20.1.^39^ Models of D2R_T4L (PDB ID: 6C38) and D2R_Fab3089 (PDB ID: 7DFP) were firstly docked into the cryo-EM map in UCSF ChimeraX-1.5^40^ followed by iterative manual adjustment in COOT0.9.8.3^41^ and real space refinement in Phenix1.19.2. The refinement statistics were provided in Supplementary Table S5. Structural figures were generated using UCSF ChimeraX-1.5.

### Computational studies

The MD protocol involved several preparation steps. Initially, protonation states of His^6.55^ and Asp^3.32^ were assessed in 1 μs simulations in three replicas each. The δ configuration of histidine and charged aspartate, which resulted in the lowest ligand RMSD values, was used in all subsequent simulations. Next, stability of the binding pose was evaluated by 4 μs unbiased MD simulations in six replicas, with apo state 4 μs simulations of the receptor in three replicas serving as a reference. These simulations were also used to verify the identity of electron densities presumed to correspond to water molecules. Subsequently, funnel metadynamics simulations were used to evaluate whether the cryo-EM pose corresponds to the actual free energy minimum on the ligand binding free energy landscape. Finally, a set of metadynamics simulations was used to investigate the basis of D_2_/D_3_ selectivity.

#### Simulation setup

The cryo-EM structure of the complex was used as a starting point in all-atom MD simulations. In unbiased simulations, the cryo-EM ligand pose was used. In metadynamics runs, the ligand was placed outside of the binding pocket in the bulk solvent. Prior to initiating the simulations, mutations present in the cryo-EM construct were reverted so that all residues matched the canonical sequence. The missing second intracellular loop was modelled using MODELLER 10.4^42^ by generating 100 models of the loop and comparing their DOPE (Discrete Optimized Protein Energy) profiles. Residues at the spot of ICL3 truncation, as well as N- and C-termini were capped with N-methyl or acetyl moieties. The receptor was immersed in a preequilibrated, complex, asymmetrical membrane bilayer with composition mimicking a physiological cell membrane.^43–49^ Neutralizing counterions were then added. The rest of the simulation box was filled with 0.15M NaCl in water. Protonation states of most residues were assigned using PROPKA 3.4.0,^24,25^, apart from Asp^3.32^ and His^6.55^. The protonation state of these two residues was established by performing all-atom unbiased simulations of the ligand-receptor complexes. Simulated systems included either δor ε state of histidine and protonated or deprotonated aspartate, and three replicas of 1 μs were performed for each system. The choice of the protonation state was made by calculating the resultant ligand RMSD and visual inspection of the trajectories. Each simulation box contained ca. 64000 atoms, with the exact number depending on the presence or absence of the ligand.

Gromacs 2023^50^ was used as a simulation engine in unbiased simulations. Metadynamics runs were performed using Gromacs 2022.3 patched with Plumed 2.8.1^51^ with the funnel metadynamics module^52^ enabled. The force fields applied were Amber03ff for protein, Stockholm lipids^53^ for the membrane, and gaff2^54^ for the ligand. Restrained Electrostatic Potential (RESP) ligand charges were obtained by the PyRED server.^55,56^ The ligand topology was built with ACPYPE v. 2022.7.21.^57^ The membrane was generated with CHARMM-GUI.^58^ The asymmetric membrane composed of eight different types of lipids, including cholesterol, was preequilibrated for 400 ns. The integration timestep was set at 2 fs. A temperature of 309.75 K was maintained by the Nose-Hoover thermostat. Pressure of 1 bar was kept by the Parrinello-Rahman barostat.

The multiple walker (5 walkers), well-tempered funnel metadynamics parameters were set as follows. The biasfactor parameter was set to 15, which corresponds to ΔT = 4336.5 K. The initial Gaussian height was set to 0.2 kJ/mol. Gaussians were added every 1 ps. RMSD walls were applied to the receptor’s Cα atoms to avoid excessive protein deformation under the influence of the additional potential.

Three collective variables (CVs) were defined to properly sample flexible ligand conformations, bias all slow degrees of freedom, and avoid significant degeneracy of states. CVs were defined as a distance between the center of mass (COM) of the ligand’s central amide and the conserved Asp^2.50^ residue along the Z axis, torsion between ligand aromatic moieties and receptor’s Asp^2.50^ and Pro^5.50^ conserved residues, and between ligand aromatic moieties and the extracellular part of TM3. Gaussian width was set to 0.05 nm, 0.43 rad, 0.43 rad, respectively. Metadynamics simulations were run for 1690 ns of simulation time per walker (5 walkers, ca. 8 μs of cumulative simulation time) until convergence was reached Supplementary Fig. S8).

In the regular well-tempered metadynamics simulations of Phe^3.28^ rotation, CV was defined as χ_1_ torsion of the residue, with ΔT = 4336.5 K, initial Gaussian height of 0.1 kJ/mol, Gaussian width of 0.1 rad, and the deposition rate of once per 2 ps. The system was simulated for ca. 117 ns per each of the 20 walkers (ca. 2300 ns of cumulative simulation time). For D_2_R, the simulation box described above was used. For the D_3_R, the simulation box was prepared analogously, using a crystal structure (PDB code: 3PBL).

## Funding

The research was funded under an OPUS grant from the National Science Center (NCN, Poland), grant number 2017/27/B/NZ7/01767 (to A.A.K). The work was supported by the Polish National Agency for Academic Exchange within the Bekker NAWA Programme, project PANALLOS, BPN/BEK/2021/1/00408/U/00001, and the Sven and Lilly Lawski Foundation, stipend no. N2024-0043 (to D.B.). Calculations were performed under PRACE European Computing Infrastructure grants MOLTRANSREC and NERVOMOLSIM (to A.A.K.), Poznań Supercomputing and Networking Center grant MOLTRANSREC (to A.A.K.), Interdisciplinary Center for Mathematical and Computational Modeling in Warsaw grant, grant number G85-948 (to A.A.K). This work was also funded by the Knut and Alice Wallenberg Foundation (KAW 2019.0130) and the Swedish Research Council (2021-4186 and 2025-06266). The MD simulations were performed using resources provided by the National Academic Infrastructure for Supercomputing in Sweden (NAISS), partially funded by the Swedish Research Council through grant agreement no. 2022-06725. M.C. acknowledges funding from the Spanish Ministry of Economy and Competitiveness (MINECO, Grant number SAF2014-57138-C2-1-R to M.C.) for *in vitro* pharmacological studies, and A.G.S. acknowledges personnel fellowship (ayudas de apoyo a la etapa predoctoral; Plan gallego de investigación, innovación y crecimiento Plan I2C) from XUNTA de Galicia. P.S. acknowledges the support of the Foundation for Polish Science (FNP) (the START program). X.L. acknowledge funding from Tsinghua-Peking Center for Life Sciences, Beijing Frontier Research Center for Biological Structure, Tsinghua University, and by Tsinghua University Initiative Scientific Research Program.

## Data availability

The cryo-EM density map for D_2_R-D2AAK2-Fab3089 has been deposited in the Electron Microscopy Data Bank (EMDB) under accession code EMD-67996. Atomic coordinates for the atomic model of D_2_R-D2AAK2-Fab3089 have been deposited in the Protein Data Bank (PDB) under the accession numbers 21TZ.

## Author contributions

**Q.S**.: Methodology; Investigation; Writing – Original Draft; Visualization.

**G.H**.: Methodology; Investigation; Writing – Original Draft; Visualization.

**D.B**.: Methodology; Formal analysis; Investigation; Writing – Original Draft; Writing – Review & Editing; Visualization; Funding acquisition.

**A.G.S**.: Methodology; Formal analysis; Investigation; Visualization; Funding acquisition.

**E.K**.: Methodology; Formal analysis; Investigation; Writing – Original Draft; Visualization.

**P.S**.: Methodology; Formal analysis; Investigation; Writing – Original Draft; Funding acquisition.

**A.A**.: Investigation.

**K.M.T.-D**.: Methodology; Formal analysis; Investigation; Writing – Original Draft; Writing – Review & Editing; Visualization; Supervision.

**T.M.W**.: Methodology; Formal analysis; Investigation; Writing – Original Draft; Writing – Review & Editing; Supervision.

**M.C**.: Methodology; Formal analysis; Resources; Writing – Original Draft; Writing – Review & Editing; Visualization; Supervision; Funding acquisition.

**J.C**.: Methodology; Resources; Writing – Original Draft; Writing – Review & Editing; Supervision; Funding acquisition.

**X.L**.: Methodology; Formal analysis; Investigation; Resources; Writing – Original Draft; Writing – Review & Editing; Visualization; Supervision; Funding acquisition.

**A.A.K**.: Conceptualization; Methodology; Formal analysis; Investigation; Resources; Writing – Original Draft; Writing – Review & Editing; Supervision; Project administration; Funding acquisition.

## Notes

https://www.rcsb.org/structure/unreleased/21TZ

### Competing Interest Statement

The authors have declared no competing interest.

## References

1. Cheng, L. et al. Structure, function and drug discovery of GPCR signaling. Mol. Biomed. 4, 46 (2023).

2. Speranza, L., Miniaci, M. C. & Volpicelli, F. The Role of Dopamine in Neurological, Psychiatric, and Metabolic Disorders and Cancer: A Complex Web of Interactions. Biomedicines 13, 492 (2025).

3. Podlewska, S. et al. Low Basicity as a Characteristic for Atypical Ligands of Serotonin Receptor 5-HT2. Int. J. Mol. Sci. 22, 1035 (2021).

4. Vandenberg, J. I. et al. hERG K(+) channels: structure, function, and clinical significance. Physiol. Rev. 92, 1393–1478 (2012).

5. Sriram, K. & Insel, P. A. G Protein-Coupled Receptors as Targets for Approved Drugs: How Many Targets and How Many Drugs? Mol. Pharmacol. 93, 251–258 (2018).

6. Hauser, A. S. et al. GPCR activation mechanisms across classes and macro/microscales. Nat. Struct. Mol. Biol. 28, 879–888 (2021).

7. Liu, S., Anderson, P. J., Rajagopal, S., Lefkowitz, R. J. & Rockman, H. A. G Protein-Coupled Receptors: A Century of Research and Discovery. Circ. Res. 135, 174–197 (2024).

8. Ballesteros, J. A., Weinstein, H. Integrated methods for the construction of three-dimensional models and computational probing of structure-function relations in G protein-coupled receptors. in Methods in Neurosciences (ed. Sealfon, S. C.) vol. 25 366–428 Academic Press, 1995.

9. Ladduwahetty, T. et al. Non-basic ligands for aminergic GPCRs: the discovery and development diaryl sulfones as selective, orally bioavailable 5-HT2A receptor antagonists for the treatment of sleep disorders. Bioorg. Med. Chem. Lett. 20, 3708–3712 (2010).

10. Hayatshahi, H. S. et al. Factors Governing Selectivity of Dopamine Receptor Binding Compounds for D2R and D3R Subtypes. J. Chem. Inf. Model. 61, 2829–2843 (2021).

11. Luedtke, R. R. et al. Comparison of the Binding and Functional Properties of Two Structurally Different D2 Dopamine Receptor Subtype Selective Compounds. ACS Chem. Neurosci. 3, 1050–1062 (2012).

12. Newman, A. H. et al. Molecular Determinants of Selectivity and Efficacy at the Dopamine D3 Receptor. J. Med. Chem. 55, 6689–6699 (2012).

13. Won, S. J., Visayas, B. R. B., Lee, K. H., Capangpangan, R. Y. & Shi, L. Decoding the Structure–Activity Relationship of the Dopamine D3 Receptor-Selective Ligands Using Machine and Deep Learning Approaches. J. Chem. Inf. Model. 65, 5495–5507 (2025).

14. Legros, C. et al. Approach to the specificity and selectivity between D2 and D3 receptors by mutagenesis and binding experiments part I: Expression and characterization of D2 and D3 receptor mutants. Protein Sci. 31, e4459 (2022).

15. Möller, D. et al. Functionally Selective Dopamine D2, D3 Receptor Partial Agonists. J. Med. Chem. 57, 4861–4875 (2014).

16. Kaczor, A. A. et al. Structure-Based Virtual Screening for Dopamine D2 Receptor Ligands as Potential Antipsychotics. ChemMedChem 11, 718–729 (2016).

17. Stępnicki, P. et al. Development and Characterization of Novel Selective, Non-Basic Dopamine D2 Receptor Antagonists for the Treatment of Schizophrenia. Molecules 28, 4211 (2023).

18. Motulsky, H.; Christopoulos, A. Fitting Models to Biological Data Using Linear and Nonlinear Regression; GraphPad Software Inc.: San Diego, 2003.

19. Steinfeld, T., Mammen, M., Smith, J. A. M., Wilson, R. D. & Jasper, J. R. A novel multivalent ligand that bridges the allosteric and orthosteric binding sites of the M2 muscarinic receptor. Mol. Pharmacol. 72, 291–302 (2007).

20. Mistry, S. N. et al. Discovery of a Novel Class of Negative Allosteric Modulator of the Dopamine D2 Receptor Through Fragmentation of a Bitopic Ligand. J. Med. Chem. 58, 6819–6843 (2015).

21. Sykes, D. A. et al. Extrapyramidal side effects of antipsychotics are linked to their association kinetics at dopamine D2 receptors. Nat. Commun. 8, 763 (2017).

22. Wang, S. et al. Structure of the D2 dopamine receptor bound to the atypical antipsychotic drug risperidone. Nature 555, 269–273 (2018).

23. Im, D. et al. Structure of the dopamine D2 receptor in complex with the antipsychotic drug spiperone. Nat. Commun. 11, 6442 (2020).

24. Olsson, M. H. M., Søndergaard, C. R., Rostkowski, M. & Jensen, J. H. PROPKA3: Consistent Treatment of Internal and Surface Residues in Empirical pKa Predictions. J. Chem. Theory Comput. 7, 525–537 (2011).

25. Søndergaard, C. R., Olsson, M. H. M., Rostkowski, M. & Jensen, J. H. Improved Treatment of Ligands and Coupling Effects in Empirical Calculation and Rationalization of pKa Values. J. Chem. Theory Comput. 7, 2284–2295 (2011).

26. Dror, R. O. et al. Pathway and mechanism of drug binding to G-protein-coupled receptors. Proc. Natl. Acad. Sci. U.S.A. 108, 13118–13123 (2011).

27. Kaczor, A. A., Żuk, J. & Matosiuk, D. Comparative molecular field analysis and molecular dynamics studies of the dopamine D2 receptor antagonists without a protonatable nitrogen atom. Med. Chem. Res. 27, 1149–1166 (2018).

28. Jakowiecki, J., Orzeł, U., Chawananon, S., Miszta, P. & Filipek, S. The Hydrophobic Ligands Entry and Exit from the GPCR Binding Site-SMD and SuMD Simulations. Molecules 25, 1930 (2020).

29. Venkatakrishnan, A. J. et al. Diverse GPCRs exhibit conserved water networks for stabilization and activation. Proc. Natl. Acad. Sci. U. S. A. 116, 3288–3293 (2019).

30. Zou, R. et al. The role of metal ions in G protein-coupled receptor signalling and drug discovery. WIREs Comput. Mol. Sci. 12, e1565 (2022).

31. Harding, M. M., Nowicki, M. W. & Walkinshaw, M. D. Metals in protein structures: a review of their principal features. Crystallogr. Rev. 16, 247–302 (2010).

32. Bellissent-Funel, M.-C. et al. Water Determines the Structure and Dynamics of Proteins. Chem. Rev. 116, 7673–7697 (2016).

33. Selvam, B., Shamsi, Z. & Shukla, D. Universality of the Sodium Ion Binding Mechanism in Class A G-Protein-Coupled Receptors. Angew. Chem. Int. Ed. 57, 3048–3053 (2018).

34. Chan, H. C. S. et al. Enhancing the Signaling of GPCRs via Orthosteric Ions. ACS Cent. Sci. 6, 274–282 (2020).

35. Seeman, P. All Roads to Schizophrenia Lead to Dopamine Supersensitivity and Elevated Dopamine D2High Receptors. CNS Neurosci. Ther. 17, 118–132 (2011).

36. Lu, C.-T. et al. Current approaches to enhance CNS delivery of drugs across the brain barriers. Int. J. Nanomedicine 9, 2241–2257 (2014).

37. Zheng, S. Q. et al. MotionCor2: anisotropic correction of beam-induced motion for improved cryo-electron microscopy. Nat. Methods 14, 331–332 (2017).

38. Punjani, A., Rubinstein, J. L., Fleet, D. J. & Brubaker, M. A. cryoSPARC: algorithms for rapid unsupervised cryo-EM structure determination. Nat. Methods 14, 290–296 (2017).

39. Liebschner, D. et al. Macromolecular structure determination using X-rays, neutrons and electrons: recent developments in Phenix. Acta Crystallogr. D Struct. Biol. 75, 861–877 (2019).

40. Pettersen, E. F. et al. UCSF ChimeraX: Structure visualization for researchers, educators, and developers. Protein Sci. 30, 70–82 (2021).

41. Emsley, P., Lohkamp, B., Scott, W. G. & Cowtan, K. Features and development of Coot. Acta Crystallogr. D Biol. Crystallogr. 66, 486–501 (2010).

42. Webb, B. & Sali, A. Comparative Protein Structure Modeling Using MODELLER. Current Protocols in Bioinformatics 54, 5.6.1-5.6.37 (2016).

43. Ingólfsson, H. I. et al. Computational Lipidomics of the Neuronal Plasma Membrane. Biophys. J. 113, 2271–2280 (2017).

44. Koldsø, H. & Sansom, M. S. P. Organization and Dynamics of Receptor Proteins in a Plasma Membrane. J. Am. Chem. Soc. 137, 14694–14704 (2015).

45. Celver, J., Sharma, M. & Kovoor, A. D(2)-Dopamine receptors target regulator of G protein signaling 9-2 to detergent-resistant membrane fractions. J. Neurochem. 120, 56–69 (2012).

46. Sharma, M., Celver, J., Octeau, J. C. & Kovoor, A. Plasma Membrane Compartmentalization of D2 Dopamine Receptors*. J. Biol. Chem. 288, 12554–12568 (2013).

47. Parton, D. L., Klingelhoefer, J. W. & Sansom, M. S. P. Aggregation of Model Membrane Proteins, Modulated by Hydrophobic Mismatch, Membrane Curvature, and Protein Class. Biophys. J. 101, 691–699 (2011).

48. Ng, H. W., Laughton, C. A. & Doughty, S. W. Molecular dynamics simulations of the adenosine A2a receptor in POPC and POPE lipid bilayers: effects of membrane on protein behavior. J. Chem. Inf. Model. 54, 573–581 (2014).

49. Sugita, M., Fujie, T., Yanagisawa, K., Ohue, M. & Akiyama, Y. Lipid Composition Is Critical for Accurate Membrane Permeability Prediction of Cyclic Peptides by Molecular Dynamics Simulations. J. Chem. Inf. Model. 62, 4549–4560 (2022).

50. Páll, S. et al. Heterogeneous parallelization and acceleration of molecular dynamics simulations in GROMACS. J. Chem. Phys. 153, 134110 (2020).

51. Bonomi, M. et al. Promoting transparency and reproducibility in enhanced molecular simulations. Nat. Methods 16, 670–673 (2019).

52. Raniolo, S. & Limongelli, V. Ligand binding free-energy calculations with funnel metadynamics. Nat. Protoc. 15, 2837–2866 (2020).

53. Grote, F. & Lyubartsev, A. P. Optimization of Slipids Force Field Parameters Describing Headgroups of Phospholipids. J. Phys. Chem. B 124, 8784–8793 (2020).

54. He, X., Man, V. H., Yang, W., Lee, T.-S. & Wang, J. A fast and high-quality charge model for the next generation general AMBER force field. J. Chem. Phys. 153, 114502 (2020).

55. Vanquelef, E. et al. R.E.D. Server: a web service for deriving RESP and ESP charges and building force field libraries for new molecules and molecular fragments. Nucleic Acids Res. 39, W511–517 (2011).

56. Dupradeau, F.-Y. et al. The R.E.D. Tools: Advances in RESP and ESP charge derivation and force field library building. Phys. Chem. Chem. Phys. 12, 7821–7839 (2010).

57. Sousa da Silva, A. W. & Vranken, W. F. ACPYPE - AnteChamber PYthon Parser interfacE. BMC Res. Notes 5, 367 (2012).

58. Jo, S., Kim, T., Iyer, V. G. & Im, W. CHARMM-GUI: a web-based graphical user interface for CHARMM. J. Comput. Chem. 29, 1859–1865 (2008).

