## Supplementary material for "Structural Dynamics of the Dopamine D_2_ Receptor with a Non-Basic Ligand": NA

**CONTENT**

**Supplementary Results:** chemistry

**NMR spectra of compounds 1a-1d and intermediates**

**Supplementary Methods:**

Chemistry

*In vitro* pharmacological studies

*In vivo* studies

**Supplementary figures**

**Fig. S1.** D2AAK2 shows high selectivity for the D_2_R.

**Fig. S2.** Lack of effect of D2AAK2 on binding dissociation kinetics of the orthosteric radioligand [^3^H]-spiperone from D_2_R.

**Fig. S3.** Cryo-EM data processing of D_2_R-T4L-D2AAK2-Fab3089 complex.

**Fig. S4.** Vertical cross sections of the ligand-binding pockets of D_2_R bound with different ligands.

**Fig. S5.** Comparison of D2AAK2-bound D_2_R structure with agonist-bound D_2_R structure.

**Fig. S6.** Water and sodium occupancy analysis.

**Fig. S7.** Effect of D2AAK2 (6.5, 12.5, 25, and 50 mg/kg, i.p.) on male mouse spontaneous locomotor activity.

**Fig. S8.** Convergence Analysis of Ligand Binding Simulations.

**Supplementary Tables**

**Table S1.** Modulation of quinpirole responses by D2AAK2 in cAMP functional data at D_2_R fits globally to the modified Gaddum/Schild EC_50_-shift model of competitive antagonism.

**Table S2.** Results for D2AAK2 derivatives 1a-d (10 µM) in competition radioligand binding assays at D_2_R.

**Table S3.** Statistical analysis of D2AAK2 effect (6.5, 12.5, and 25 mg/kg) on amphetamine-induced animal hyperactivity.

**Table S4.** Detailed experimental conditions employed in competition radioligand binding assays.

**Table S5.** Cryo-EM data collection, reﬁnement, and validation statistics.

**References**

**Supplementary Results**

Chemistry

All final compounds were synthesized starting from the common 4-trifluoromethylaniline. This starting material was acylated with 3-chloropropionyl chloride, affording 3-chloro-*N*-(4-(trifluoromethyl)phenyl)propanamide in excellent yield, which in turn was used to alkylate pyrrolidine, piperidine, morpholine, and 2*H*-benzo[*b*][1,4]thiazin-3(4*H*)-one affording compounds **1a**, **1b, 1c,** and **1d,** respectively (Scheme 1). Compound **1e** was approached similarly. In this case, we found it was more practical to methylate starting aniline first and then conduct subsequent acylation and alkylation analogous to the first four compounds (Scheme 2).

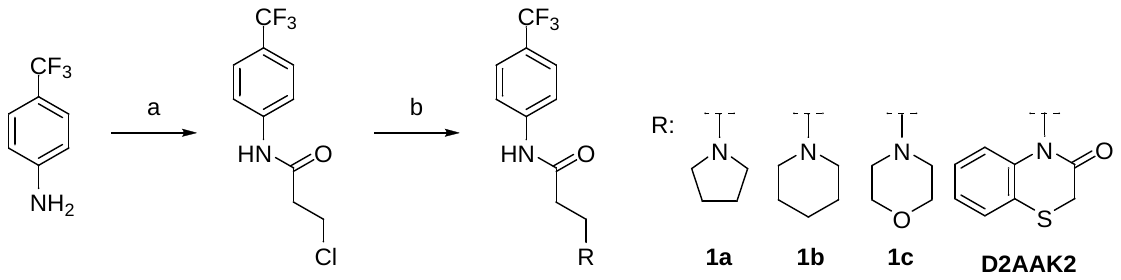

**Scheme** 1. Synthesis of compounds **1a**-**d**. Reagents and conditions: a) 3-chloropropionyl chloride, toluene, reflux, 2h, 92%; b) amine, K_2_CO_3_, DMF, rt, 4h, 4-15% (for **1a**-**c**); lactam, K_2_CO_3_, ACN, reflux, 24h, 34% (for **D2AAK2**).

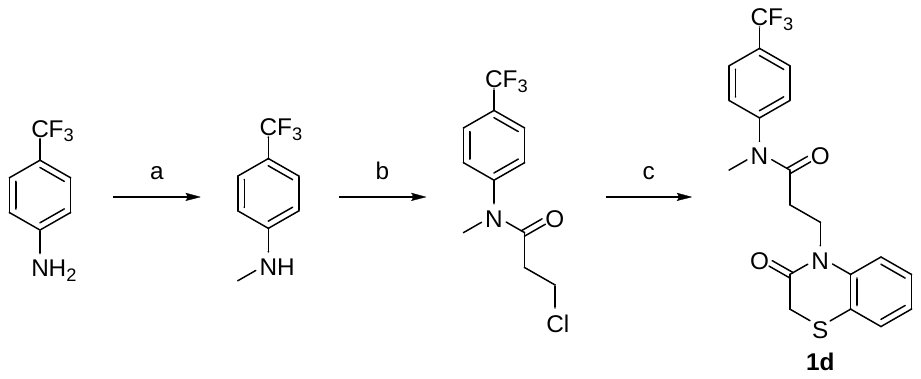

**Scheme 2.** Synthesis of compound 1d. Reagents and conditions: a) paraformaldehyde, MeONa, MeOH, reflux, 2h, then NaBH4, 0 °C to reflux, 2h, 82%; b) 3-chloropropionyl chloride, K_2_CO_3_, H_2_O/acetone; 0 to 30 °C, 24h, 94%; c) 2H-benzo[b][1,4]thiazin-3(4H)-one, K_2_CO_3_, KI, MeCN, reflux, 24h, 31%

***N*-methyl-4-(trifluoromethyl)aniline**

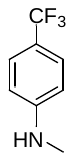

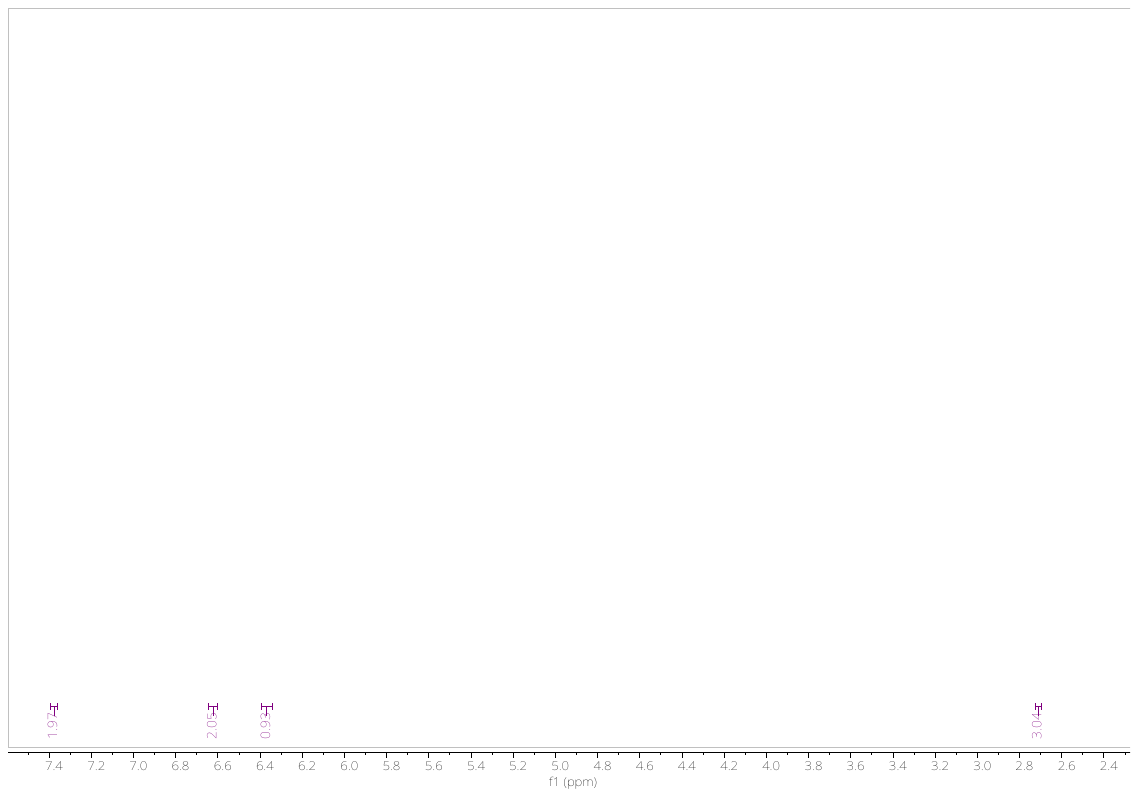

**3-chloro-*N*-methyl-*N*-(4-(trifluoromethyl)phenyl)propanamide**

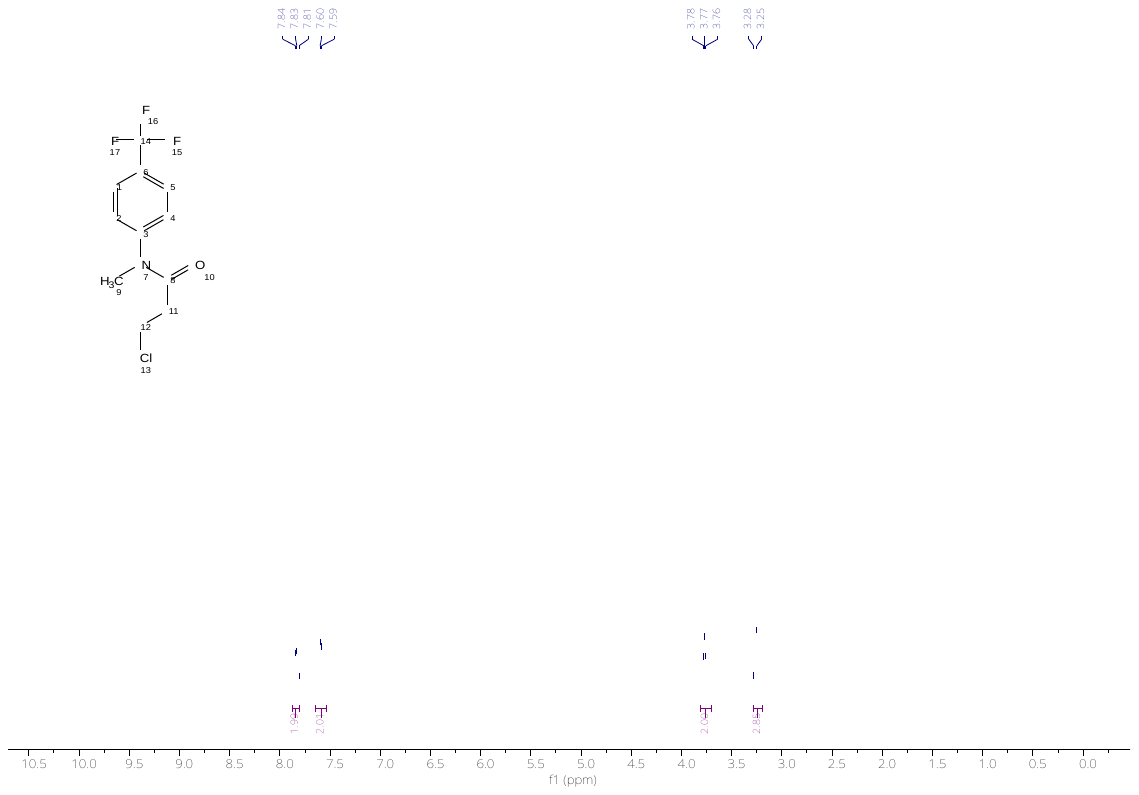

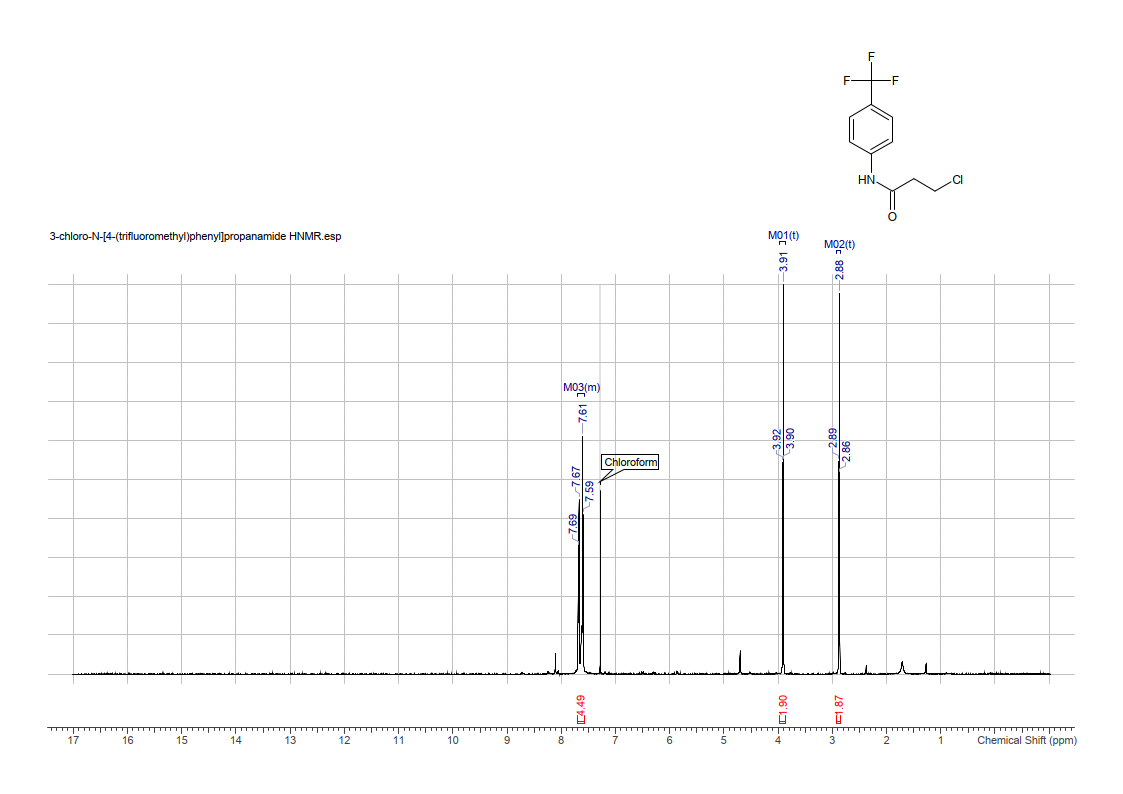

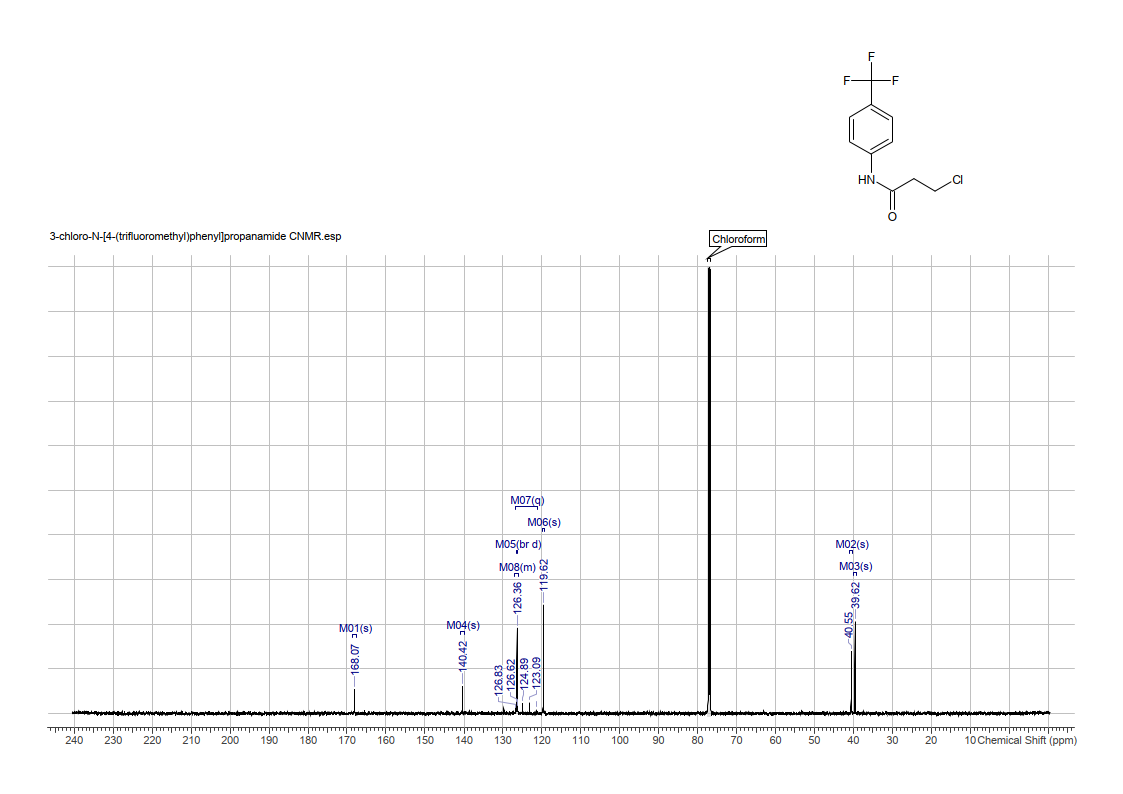

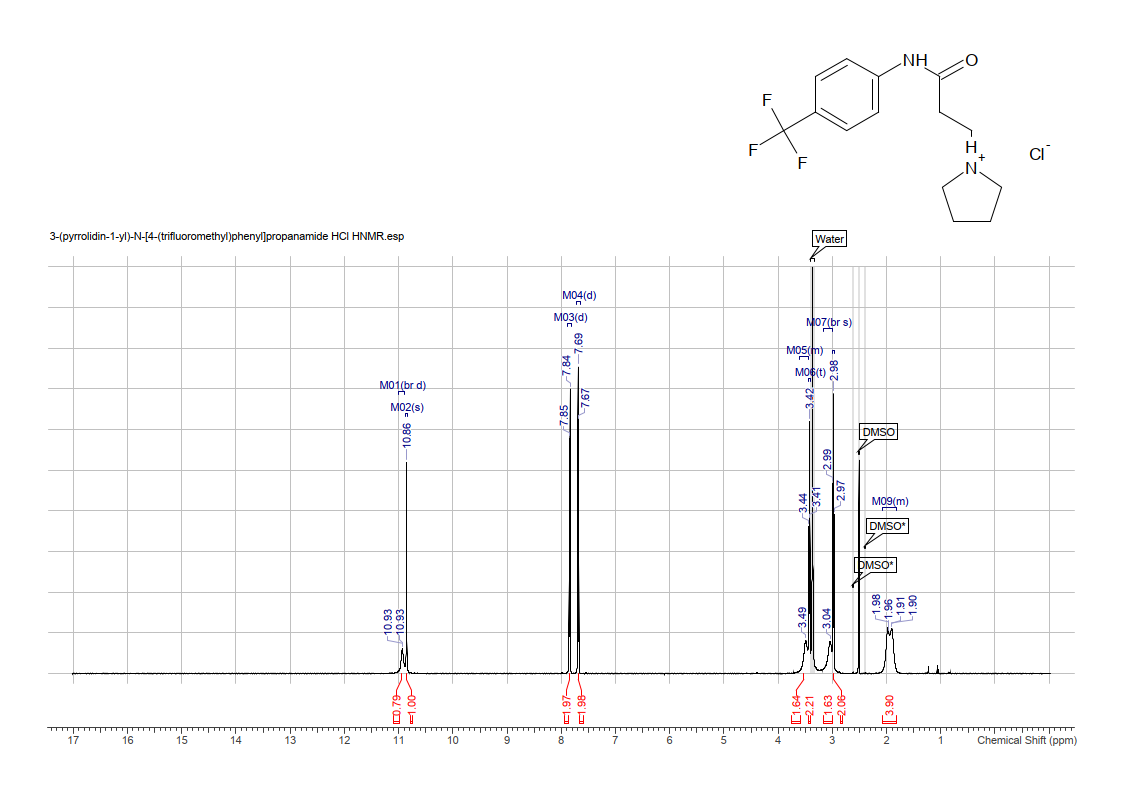

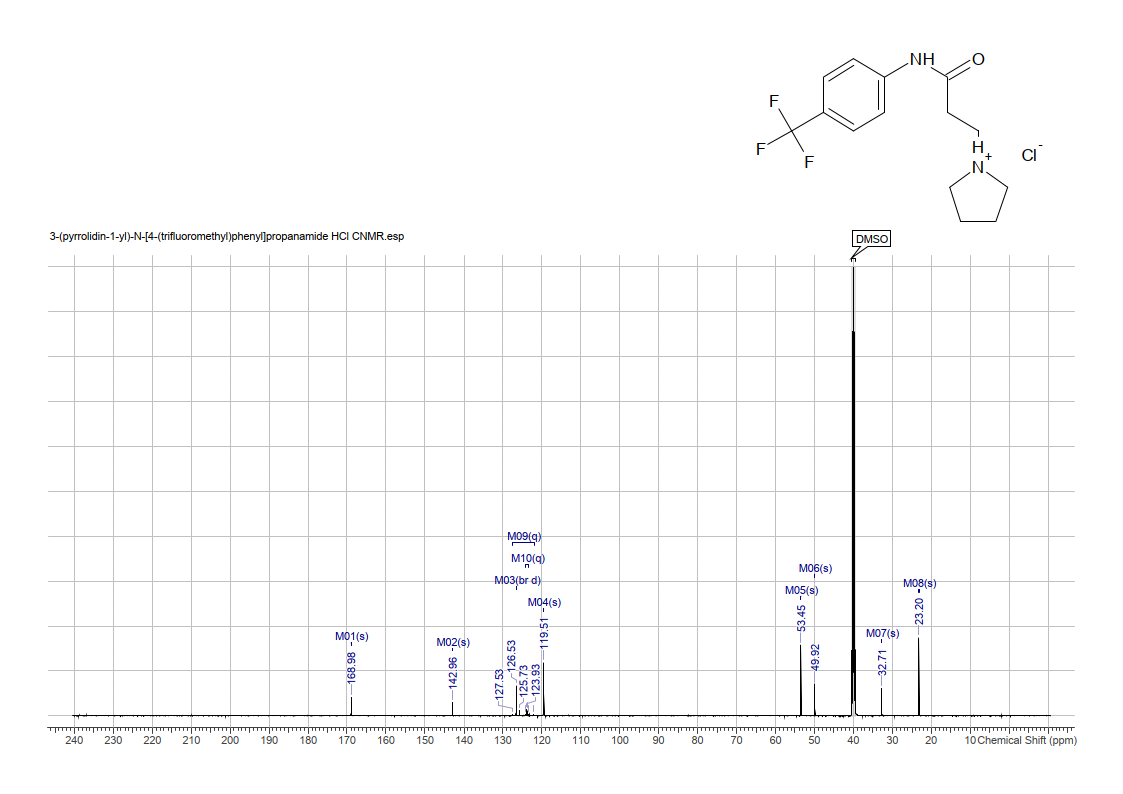

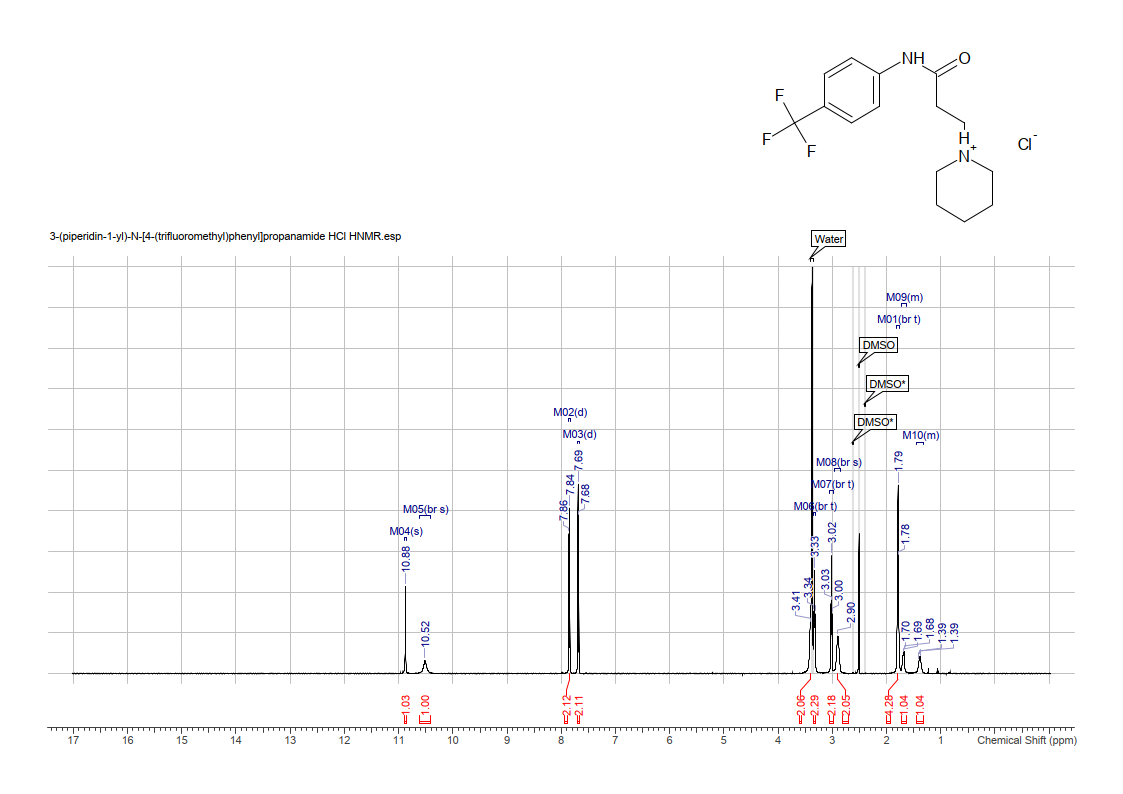

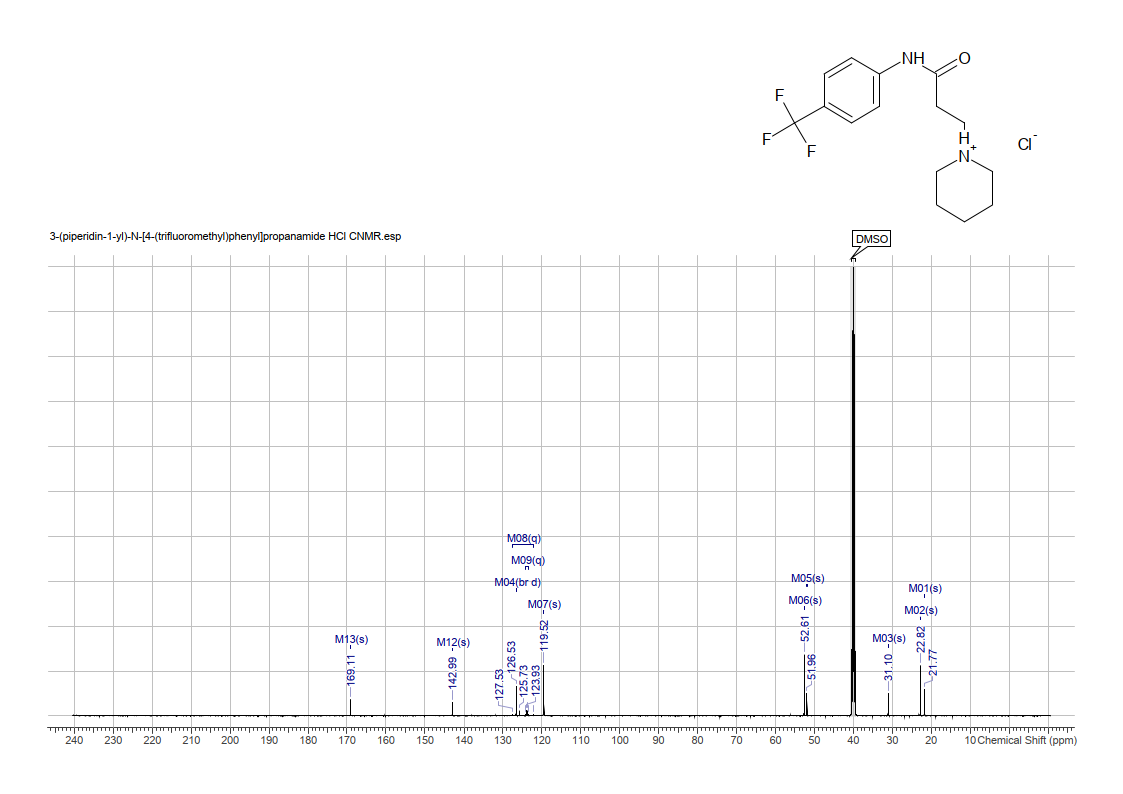

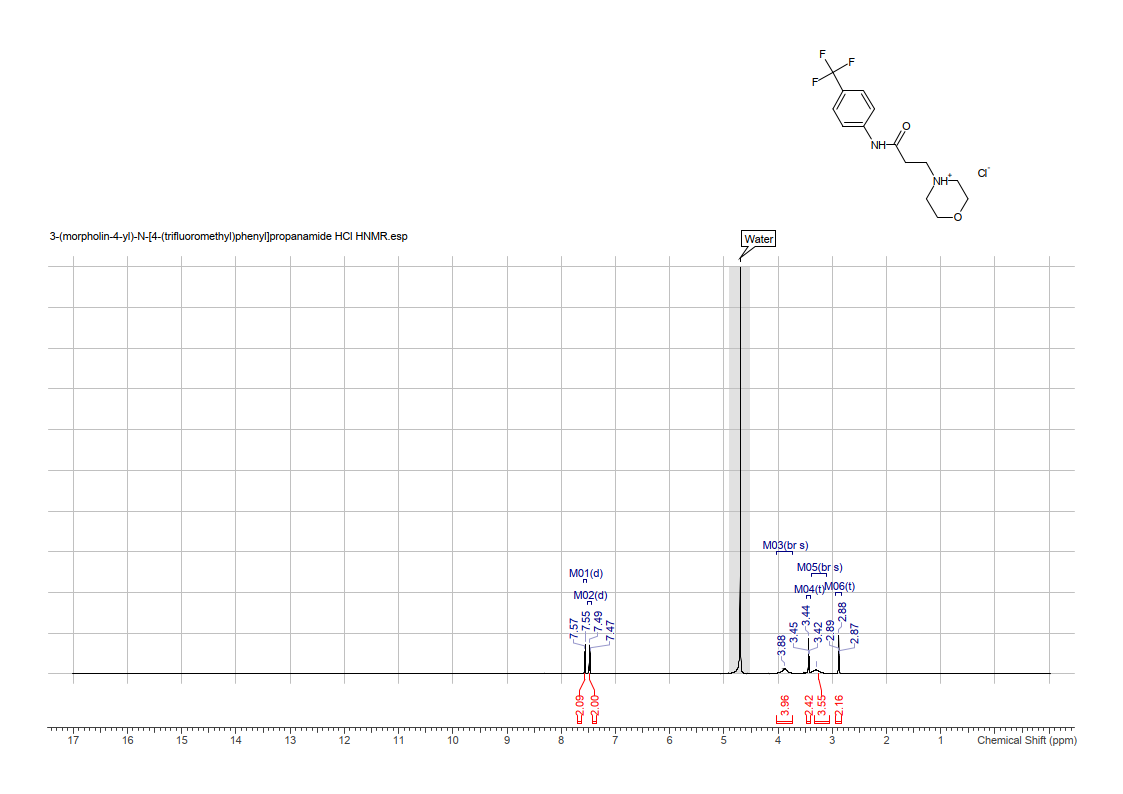

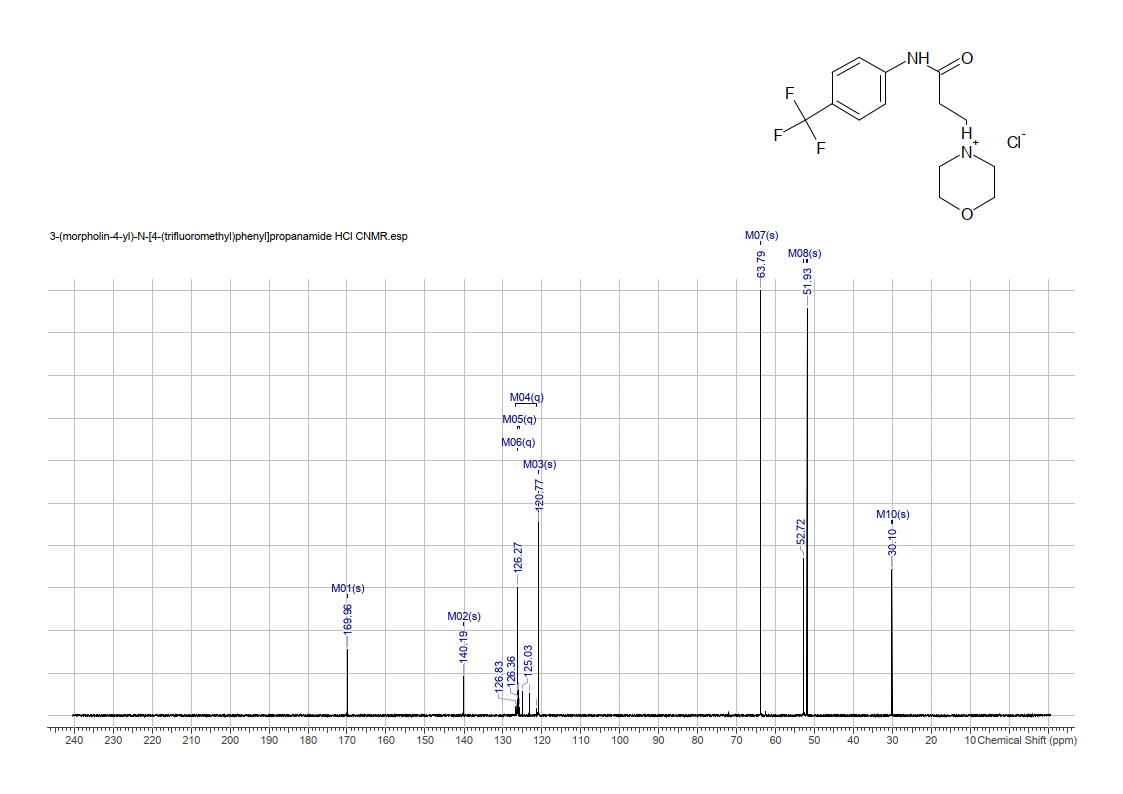

**3-(3-oxo-2,3-dihydro-4*H*-benzo[*b*][1,4]thiazin-4-yl)-*N*-(4-(trifluoromethyl)phenyl)propanamide (D2AAK2)**
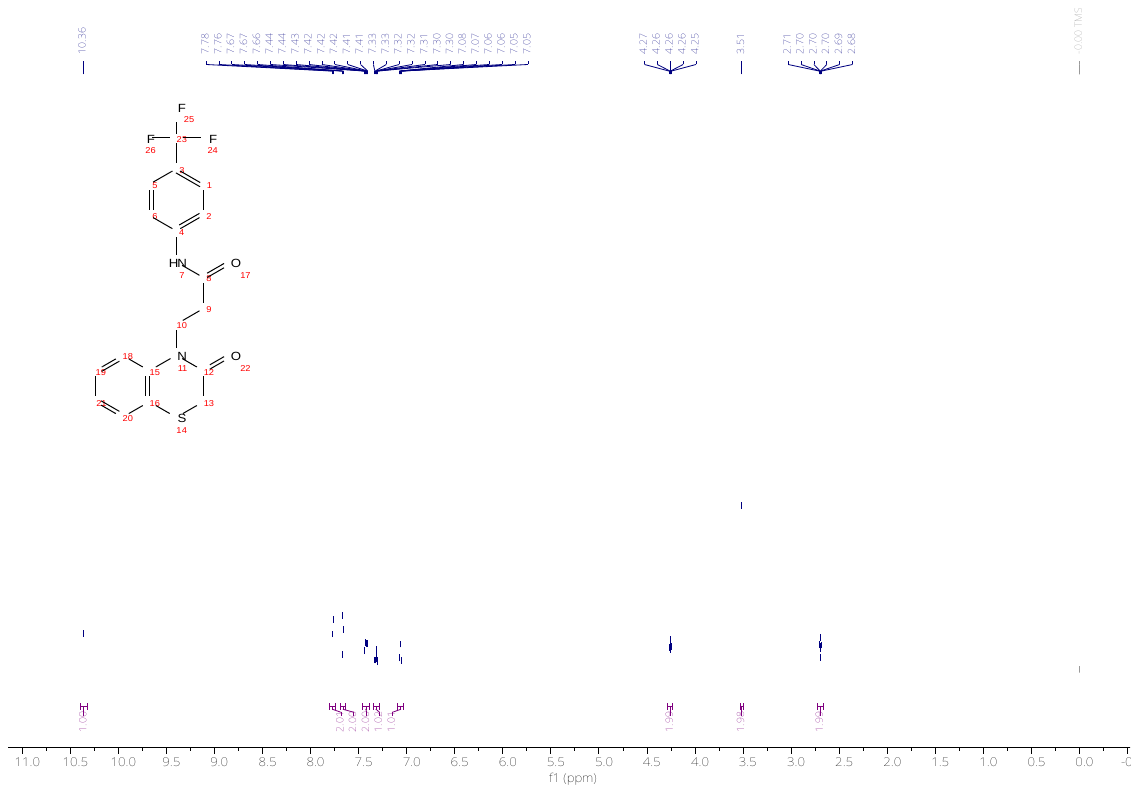

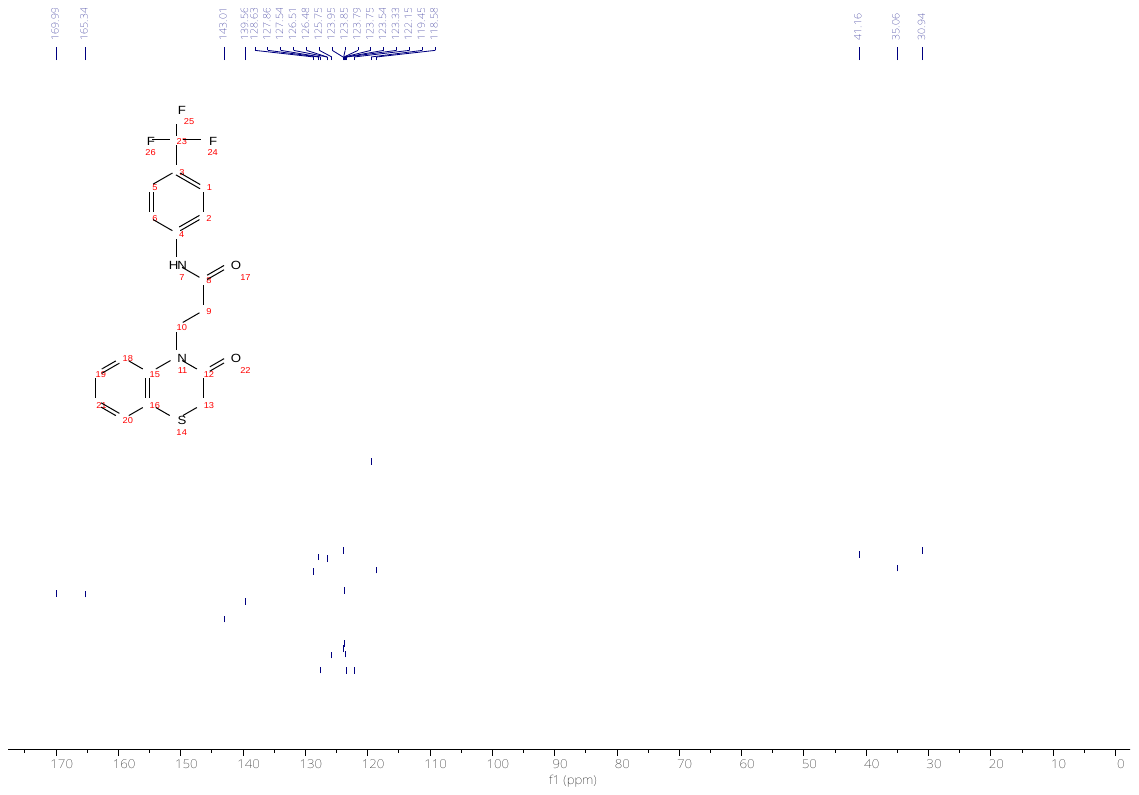

***N*-methyl-3-(3-oxo-2,3-dihydro-4*H*-benzo[*b*][1,4]thiazin-4-yl)-*N*-(4-(trifluoromethyl)phenyl)propanamide 1d**

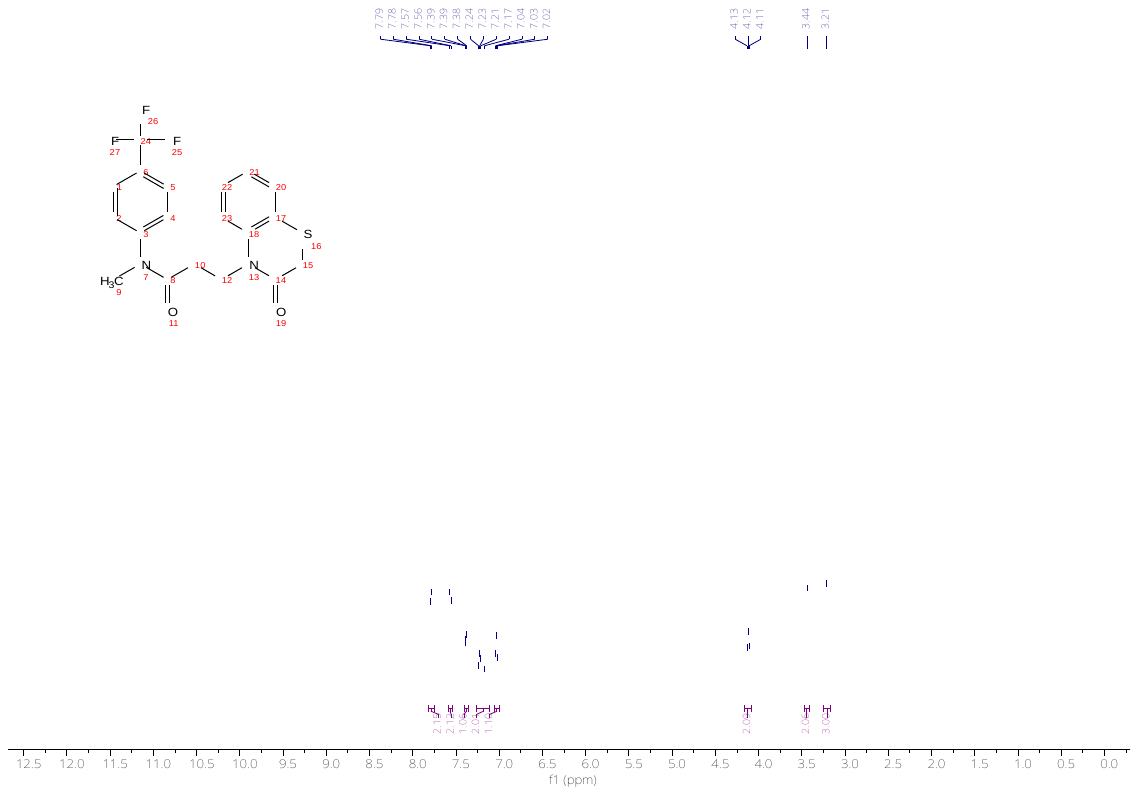

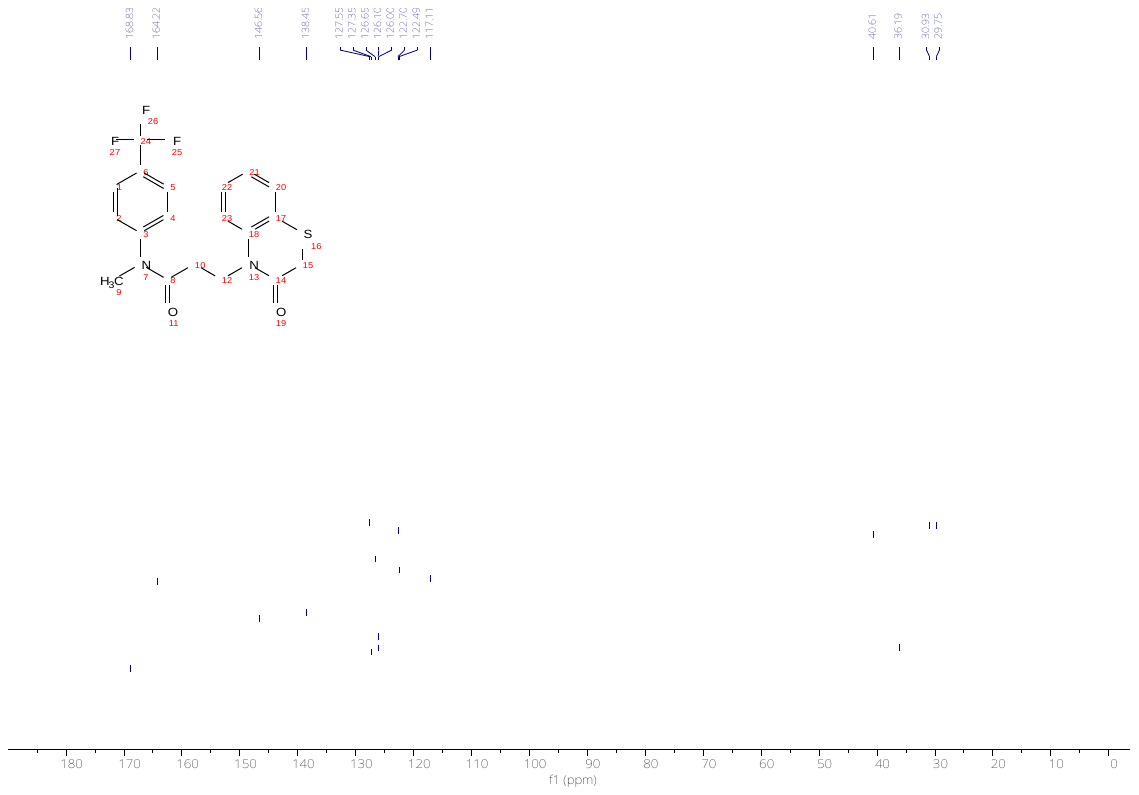

**Supplementary Methods**

**Chemistry**

**3-chloro-*N*-(4-(trifluoromethyl)phenyl)propanamide**
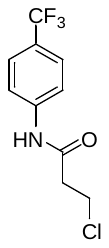

A solution of 3-chloropropionyl chloride (16.6 g, 0.131 mol) in 50 mL of toluene was added dropwise to 4-trifluoromethylaniline (19.9 g, 0.124 mol) dissolved in 200 mL of toluene dropwise over the course of 20 minutes. The reaction was refluxed for 2 hours and then cooled in an ice-water bath. The product was filtered and washed with cold toluene (75 mL) to afford a white solid (28.3 g, 91%).

^1^H NMR (600MHz, CHLOROFORM-d) δ = 7.71 - 7.58 (m, 4H), 3.91 (t, *J*=6.3 Hz, 2H), 2.88 (t, *J*=6.4 Hz, 2H)
^13^C NMR (151MHz, CHLOROFORM-d) δ = 168.1, 140.4, 126.4 (br d, *J*=3.5 Hz), 126.9 - 126.1 (m), 124.0 (q, *J*=271.8 Hz), 119.6, 40.6, 39.6

MS [M+H]^+^ teor m/z = 252 exp m/z = 252

**General procedure to obtain compounds 1a-c**

A flask was charged with 3-chloro-*N*-(4-(trifluoromethyl)phenyl)propanamide (1.26 g, 5 mmol), potassium carbonate (1.38 g, 10 mmol), amine (5 mmol), and DMF. The reaction was stirred at room temperature for the prescribed time. Solvent was removed on a rotavap, and the remaining residue was taken up in 30 mL of DCM. It was then extracted three times with 5% HCl, made alkaline with 5% NaOH, and extracted with DCM. After solvent removal, the remaining residue was taken up in ether. After the addition of anhydrous HCl in ethanol, a precipitate formed, which was filtered and washed.

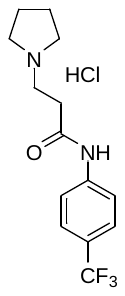
**3-(pyrrolidin-1-yl)-*N*-(4-(trifluoromethyl)phenyl)propanamide hydrochloride 1a**

Reaction time 4 hours using 50 mL of DMF. Product was obtained as a white solid (235 mg, 15%) in the form of a hydrochloride salt and characterized as such.

^1^H NMR (600MHz, DMSO-d6) δ = 10.93 (br d, *J*=2.1 Hz, 1H), 10.86 (s, 1H), 7.84 (d, *J*=8.8 Hz, 2H), 7.68 (d, *J*=8.9 Hz, 2H), 3.61 - 3.45 (m, 2H), 3.42 (t, *J*=7.5 Hz, 2H), 3.04 (br s, 2H), 2.98 (t, *J*=7.5 Hz, 2H), 2.08 - 1.82 (m, 4H)
^13^C NMR (151MHz, DMSO-d6) δ = 169.0, 143.0, 126.5 (br d, *J*=3.5 Hz), 123.8 (q, *J*=32.4 Hz), 124.8 (q, J=271.3 Hz), 119.5, 53.5, 49.9, 32.7, 23.2

MS [M+H]^+^ teor m/z = 287 exp m/z = 287

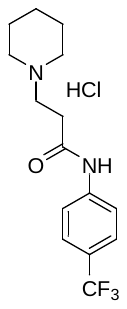
**3-(piperidin-1-yl)-N-(4-(trifluoromethyl)phenyl)propanamide hydrochloride 1b**

Reaction time 5 hours using 25 mL of DMF. Product was obtained as a white solid (69 mg, 4%) in the form of a hydrochloride salt and characterized as such.

^1^H NMR (600MHz, DMSO-d6) δ = 10.88 (s, 1H), 10.51 (br s, 1H), 7.85 (d, *J*=8.6 Hz, 2H), 7.68 (d, *J*=8.6 Hz, 2H), 3.33 (br t, *J*=7.3 Hz, 3H), 3.02 (br t, *J*=7.3 Hz, 2H), 2.90 (br s, 2H), 1.79 (br t, *J*=5.2 Hz, 5H), 1.73 - 1.64 (m, 1H), 1.45 - 1.32 (m, 1H)
^13^C NMR (151MHz, DMSO-d6) δ = 169.1, 143.0, 126.5 (br d, *J*=3.5 Hz), 123.8 (q, *J*=31.8 Hz), 124.8 (q, *J*=271.1 Hz), 119.5, 52.6, 52.0, 31.1, 22.8, 21.8

MS [M+H]^+^ teor m/z = 301 exp m/z = 301

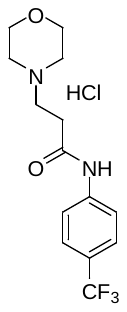
**3-morpholino-N-(4-(trifluoromethyl)phenyl)propanamide hydrochloride 1c**

Reaction time 6 hours using 10 mL of DMF. Product was obtained as a white solid (106 mg, 6%) in the form of a hydrochloride salt and characterized as such.

^1^H NMR (600MHz, DEUTERIUM OXIDE) δ = 7.56 (d, *J*=8.6 Hz, 2H), 7.48 (d, *J*=8.4 Hz, 2H), 3.88 (br s, 4H), 3.44 (t, *J*=6.9 Hz, 2H), 3.30 (br s, 4H), 2.88 (t, *J*=6.9 Hz, 2H)
^13^C NMR (151MHz, DEUTERIUM OXIDE) δ = 170.0, 140.2, 126.3 (q, *J*=3.9 Hz), 126.0 (q, J=32.6 Hz), 124.1 (q, *J*=271.1 Hz), 120.8, 63.8, 52.7, 51.9, 30.1

MS [M+H]+ teor m/z = 303 exp m/z = 303

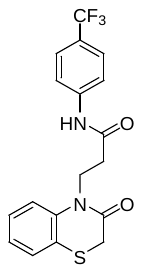
**3-(3-oxo-2,3-dihydro-4*H*-benzo[*b*][1,4]thiazin-4-yl)-*N*-(4-(trifluoromethyl)phenyl)propanamide (D2AAK2)**

A flask was charged with 3-chloro-*N*-(4-(trifluoromethyl)phenyl)propanamide (0.755 g, 3 mmol), 2*H*‑benzo[*b*][1,4]thiazin-3(4*H*)-one (0.496 g, 3 mmol), potassium carbonate (0.829 g, 6 mmol), and acetonitrile (12 mL). The reaction mixture was refluxed for 24 h, after which it was allowed to cool to ambient temperature. The solids were filtered off and the solvent was evaporated. The obtained crude product was purified by crystallization from methanol, giving the final compound as a white crystalline solid (0.391 g, 34%).

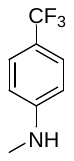
***N*-methyl-4-(trifluoromethyl)aniline**

The compound was synthesized according to the previously reported procedure [1]. Sodium methoxide was prepared by adding slowly sodium metal (460 mg, 20 mmol) to methanol (40 mL) at 0 °C. To obtained solution, 4-(trifluoromethyl)aniline (645 mg, 4 mmol) and paraformaldehyde (300 mg, 10 mmol) were added at room temperature, and then the reaction mixture was refluxed for 2 h. To obtained mixture containing imine intermediate, sodium borohydride (227 mg, 6 mmol) was added at 0 °C, and then it was refluxed for 2 h. The reaction mixture was cooled to room temperature, and the solvent was evaporated. The obtained residue was diluted with dichloromethane (30 mL), washed with water (30 mL), and dried over anhydrous sodium sulfate. The solvent was evaporated to give the product as a yellow oil (0.578 g, 82%), which was used in the next step without further purification.

^1^H NMR (600 MHz, DMSO) δ 7.38 (d, *J* = 8.7 Hz, 2H), 6.62 (d, *J* = 8.7 Hz, 2H), 6.37 (q, *J* = 5.0 Hz, 1H), 2.71 (d, *J* = 5.0 Hz, 3H).

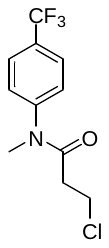
**3-chloro-*N*-methyl-*N*-(4-(trifluoromethyl)phenyl)propanamide**

The compound was obtained by modifying the previously described procedure [2]. To a stirred mixture of *N*-methyl-4-(trifluoromethyl)aniline (280 mg, 1.6 mmol), potassium carbonate (442 mg, 3.2 mmol), water (3 mL), and acetone (1.5 mL), there was added 3-chloropropionyl chloride (305 mg, 2.4 mmol) dropwise at 0 °C. Then the reaction mixture was stirred at 30 °C overnight. After cooling to room temperature, it was transferred to ice-cold water (5 mL), and then it was extracted with dichloromethane (3×10 mL). The organic layers were combined and dried with anhydrous sodium sulfate. Upon solvent removal, there was yellow oil (0.4 g, 94%), which was used in the last step without further purification.

***N*-methyl-3-(3-oxo-2,3-dihydro-4*H*-benzo[*b*][1,4]thiazin-4-yl)-*N*-(4-(trifluoromethyl)phenyl)propanamide**
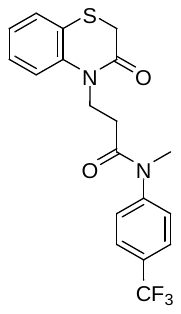
 **1d**

A reaction vial was charged with 3-chloro-*N*-methyl-*N*-(4-(trifluoromethyl)phenyl)propanamide (0.398 g, 1.5 mmol), 2*H*‑benzo[*b*][1,4]thiazin-3(4*H*)-one (0.248 g, 1.5 mmol), potassium carbonate (0.415 g, 3 mmol), catalytic amount of potassium iodide (10 mg) and acetonitrile (5 mL). The reaction mixture was refluxed overnight. Next, it was allowed to cool to room temperature, filtered, and then the solvent was evaporated. The obtained residue was purified by column chromatography (Silica gel, eluent: hexane/ethyl acetate = 1:1) giving the desired product as a colorless oil (0.181 g, 31%).

***In vitro* pharmacological studies**

***Radioligand binding assays***

Competition radioligand binding assays were performed on cell membrane preparations from cell lines stably expressing the human cloned receptors as previously reported.^1^ Chinese hamster ovary K1 (CHO-K1) cell lines stably expressing human D_2S_, 5-HT_2A_ or H_1_ receptors and human embryonic kidney 293 (HEK293) cell lines stably expressing human 5-HT_1A_ or 5-HT_7_ receptors were in-house available, whereas CHO-K1 cell lines stably expressing human D_1_ or D_3_ receptors (Perkin Elmer, Waltham, MA, USA) and Chem-1 cell line stably expressing human M_1_ receptor (Millipore) were commercially available. Detailed experimental conditions employed for each receptor are provided in Supplementary Table S4. Reactions were terminated by rapid filtration through glass fiber filters, which were washed with ice-cold wash buffer and let dry before radioactivity was quantified in a MicroBeta² microplate liquid scintillation counter (Perkin Elmer, Waltham, MA, USA). Competition binding curves of compound D2AAK2 at the different receptors were constructed using six or seven different concentrations of compound, from 0.1 nM or 1 nM till full displacement or till 30 µM or 100 µM concentrations were reached.

A possible effect of D2AAK2 on the dissociation kinetics of the orthosteric radioligand [^3^H]-spiperone from D_2_R was investigated in radioligand binding kinetic assays by an “isotopic dilution”^2^ approach, following a protocol previously described.^3^ 0.2 nM [^3^H]-spiperone was allowed to equilibrate with D_2_R cell membrane preparations for 2 hours at 30ºC in assay buffer prior to the addition of 10 µM sulpiride as a dissociator, either alone or together with D2AAK2 (10 µM), and dissociation was carried out at different time points for 180 min. Nonspecific binding was assessed in the presence of 10 µM sulpiride. Other experimental conditions, filtration and radioactivity counting were conducted as for competition radioligand binding assays.

***cAMP functional assays***

Functional assays of cAMP signaling were performed employing the same CHO-K1 cell line expressing the human D_2S_ receptor used in radioligand binding assays, and following a protocol previously described.^4^ Specifically, concentration (from 10^-11^ to 10^-3^ M)-response curves of quinpirole were performed in the absence and presence of three different concentrations (30 nM, 300 nM and 3 µM) of D2AAK2. Frozen cell stocks were thawed, resuspended in prewarmed (37 ºC) assay medium (Opti-MEM™ I medium (GIBCO, #11058021) supplemented with 500 µM 3-isobutyl-1-methylxanthine (IBMX)), and plated at a density of 1500 or 5000 cells/well, respectively in 96- (Isoplate #6005030, PerkinElmer) or 384- (#CLS3570, Corning) well microplates containing the corresponding working concentrations of D2AAK2 or assay medium. Cells were incubated for 5 min at 37ºC with gentle agitation before quinpirole was added. After 10 min incubation, forskolin (FSK) (10 uM final concentration) was added and incubation continued for 5 min. After this time, cAMP levels were quantified using homogeneous time-resolved fluorescence (HTRF)-based cAMP dynamic kit (Cisbio, Bioassays, Codolet, France) according to the manufacturer’s protocol. Basal cAMP levels were determined in the absence of D2AAK2, quinpirole, and forskolin. FSK response was determined in the absence of D2AAK2 and quinpirole. After subtraction of average intraplate basal values, quinpirole response was expressed as % of inhibition of FSK-stimulated cAMP production.

***Data Analysis***

GraphPad Prism (versions 7 and 10; San Diego, CA) was used for nonlinear regression and statistical analyses.

For competition radioligand binding assays, specific binding data were fitted to a one-site competition model:

$$Y=\text{Bottom}+\frac{\text{Top}-\text{Bottom}}{1+{10}^{(X-\log EC_{50})}}$$

where *X* is the logarithm of the molar concentration of competitor, *Y* is specific binding, and *Top* and *Bottom* represent the upper and lower plateaus of the binding curve.

The relation between competitor affinity (equilibrium dissociation constant (*K*_i_); p*K*_i_ (-log *K*_i_)) was described as:

$$\log EC_{50}=\log\left( {10}^{\log K_{i}}\left( 1+\frac{\left[ \text{Hot} \right]}{K_{d,\text{Hot}}} \right) \right)$$

were concentration of radioligand [Hot] and its affinity (*K*_d,Hot_), as determined in saturation binding experiments, were constrained to constant values.

For radioligand binding kinetic experiments, specific binding data were fitted to a one-phase exponential decay model (Y = Span · e^-^*^kX^* + Plateau), where plateau was fixed to a constant value of zero. A global fit of the data sets with shared dissociation rate constant *k* was compared with an unconstrained model in which this parameter was allowed to vary across datasets.

For cAMP functional assays, quinpirole concentration-response curves were first fitted to a four-parameter logistic equation:

$$Response= \text{Bottom}+\frac{\text{Top}-B\text{ottom}}{1+{10}^{\left( \log EC_{50}-X \right)n_{H}}}$$

where *Bottom* and *Top* represent the lower and upper asymptotes, *X* is the logarithm of agonist concentration, *log EC_50_* is the log agonist concentration that elicits 50% of the maximal response, and n*_H_* is the Hill slope. A global fit of all data sets (control and D2AAK2 at 30 nM, 300 nM and 3 µM; n = 3) to a family of curves sharing both *Top* and n*_H_* was compared with an unconstrained model in which these parameters were allowed to vary across curves. Because quinpirole-D2AAK2 interactions fulfilled the criteria for competitive antagonism at the tested D2AAK2 concentrations, the data were subsequently fitted globally to the modified logistic Gaddum/Schild EC_50_-shift model of competitive antagonism:^5^

$$Response =Bottom+ \frac{Top - Bottom}{1 + \left( \frac{{10}^{-pEC_{50}}\left( 1 + \left( \frac{\left[ B \right]}{{10}^{-pA_{2}}} \right)^{s} \right)}{\left[ A \right]} \right)^{n_{H}}}$$

where [*A*] is the agonist (quinpirole) concentration, [*B*] is antagonist (D2AAK2) concentration, *pEC_50_* is the negative logarithm of the agonist EC_50_ in the absence of antagonist, *s* is the Schild slope, *n_H_* is the shared Hill slope, and *pA_2_* is the negative logarithm of the antagonist concentration producing a twofold shift in EC_50_. Because the estimate of *s* was not significantly different from 1, the Schild slope was subsequently constrained to unity to obtain the equilibrium dissociation constant (*K*_b_), with p*A*_2_ = p*K*_b_.

Statistical comparisons between nested models were performed using the extra sum-of-squares F-test (α = 0.05). Model ranking additionally considered the Akaike’s Information Criterion corrected for small sample size (AICc)) was evaluated. Comparisons between non-nested models were based solely on AICc.

**In vivo studies**

***Drugs***

Compound D2AAK2, with a purity of 99%, synthesized as described above, was dissolved in DMSO to a final concentration of 2%, and then further diluted with an aqueous solution of 0.5% methylcellulose (MC). Both the drug and its respective vehicles were freshly prepared before each experiment and were injected intraperitoneally (i.p.). Compound D2AAK2, as well as its respective vehicles, were injected at the same volume (10 ml/kg). The purity of the compound was assessed by NMR spectral analysis.

***Animals***

The experiments were conducted using Swiss male mice, aged 2 months, obtained from the Farm of Laboratory Animals in Warszawa, Poland. These mice weighed between 20 to 30 grams. All experiments adhered to the guidelines set forth by the National Institute of Health for the Care and Use of Laboratory Animals, as well as the European Community Council Directive for Care and Use of Laboratory Animals (2010/63/*EU*). The research protocol was approved by the local ethics committee (license 55/2022), as previously documented in Kaczor et al..^6^ The mice were housed under standard laboratory conditions and were acclimated to the laboratory environment, as detailed in previous work by Kaczor et al..^6^

***Mouse spontaneous and amphetamine-induced locomotor activity***

The impact of D2AAK2 on the spontaneous locomotor activity of mice was assessed using an Opto-Varimex-4 Auto-Track animal activity meter from Columbus Instruments, USA. This evaluation took place in a sound-attenuated experimental room. The Auto-Track System relies on a grid of infrared photocells to monitor the movements of the animals. Specifically, we analyzed the arithmetic average of the distance (cm) traveled by each mouse (±SEM) in various experimental groups. In one set of experiments, each mouse was placed in a cage for 60 minutes immediately after receiving D2AAK2 (6.5, 12.5, 25, and 50 mg/kg; i.p.), amphetamine (5 mg/kg; s.c.), or a control vehicle injection. This was done to assess the effects of D2AAK2 on spontaneous locomotor activity. In another set of experiments to determine whether D2AAK2 influenced amphetamine-induced hyperactivity, each mouse received D2AAK2 (6.5, 12.5 and 25 mg/kg; i.p.) or a vehicle injection, followed by amphetamine (5 mg/kg; s.c.) or a vehicle injection 30 minutes later. Subsequently, the spontaneous locomotor activity of the mice was recorded for 30 minutes.

**Statistical analysis**

Statistical analysis was carried out using GraphPad Prism v. 8.0. One-way analysis of variance (ANOVA), followed by Bonferroni's multiple comparisons test was used to compare differences between the drug-treated groups and the control group (locomotor activity evaluations). Two-way ANOVA followed by Tukey’s post hoc test was performed after assessments of compound effects on amphetamine-induced locomotor hyperactivity. Data was expressed as mean ± SEM. A p-value < 0.05 was considered statistically significant.

**Supplementary figures**

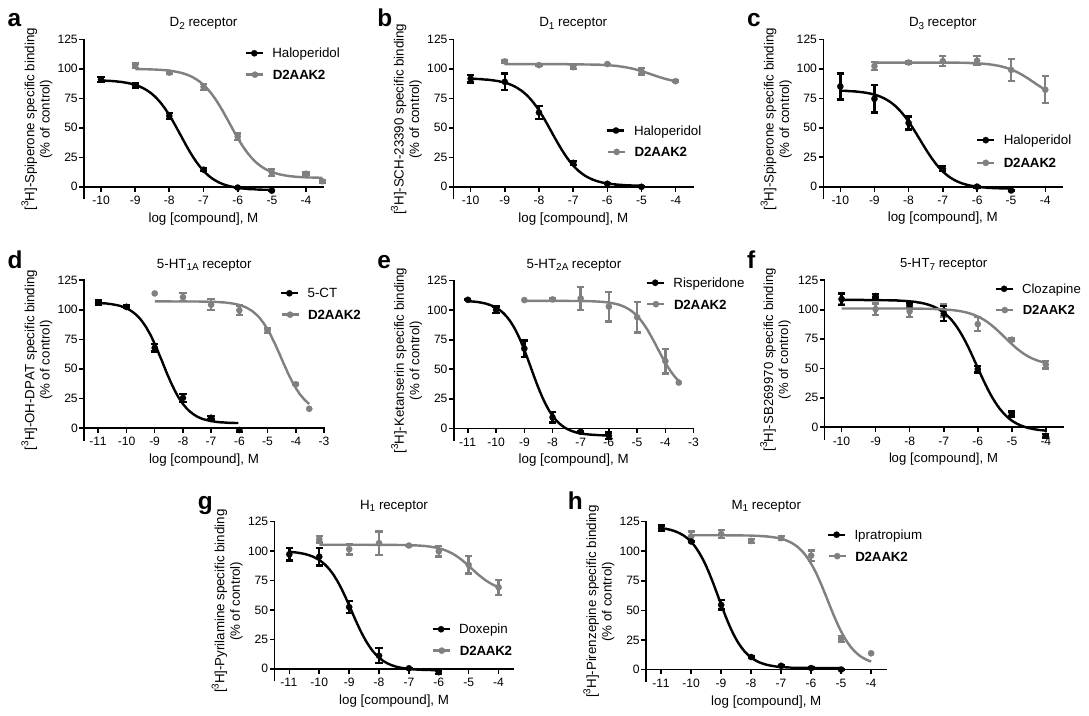

**Fig. S1. D2AAK2 shows high selectivity for the D_2_R. a-h,** Radioligand binding displacement curves for D2AAK2 and the corresponding reference compound at the indicated receptors. Graphs show data (mean ± SEM) of n = 3 (**c**, D_3_) or n = 2 independent experiments performed in duplicate.

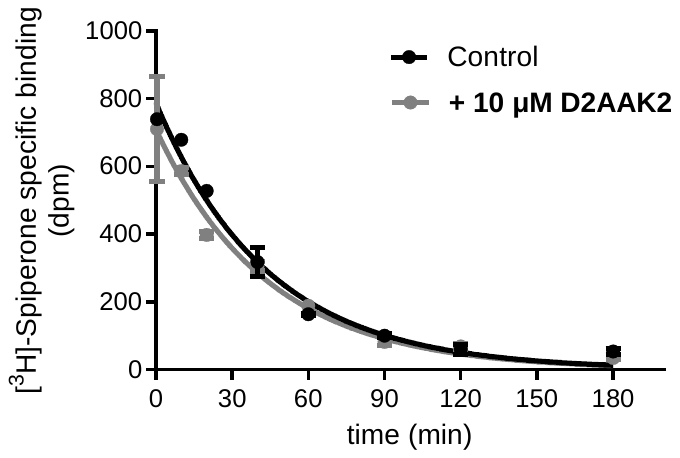

**Fig. S2. Lack of effect of D2AAK2 on binding dissociation kinetics of the orthosteric radioligand [^3^H]-spiperone from D_2_R.** Dissociation was initiated by adding dissociator either alone (control) or together with 10 µM D2AAK2, at different timepoints after equilibration of the radioligand with D_2_R membrane preparations. Data points represent the mean ± SEM of a single experiment performed in duplicate. Curves drawn through the data points represent fit to a one-phase exponential decay model with *k* value allowed to vary across datasets. Dissociation rate constant (*k*_off_) of [^3^H]-spiperone was (mean ± SEM, n = 1) 0.0229 (95% CI 0.0197–0.0264) and 0,0226 (95% CI 0.01743– 0.02924) min^-1^ for control and + 10 µM D2AAK2, respectively.

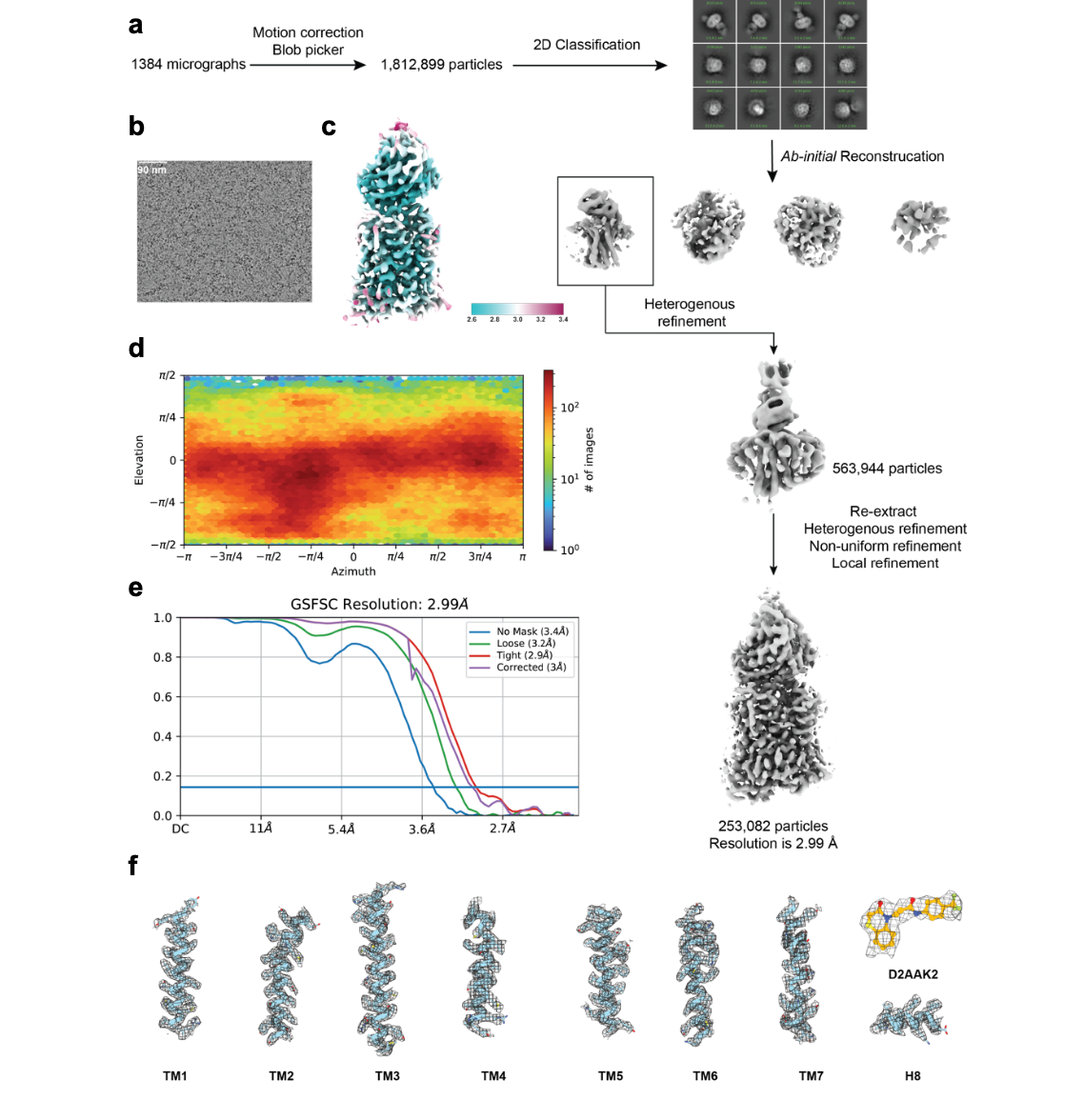

**Fig. S3. Cryo-EM data processing of D_2_R-T4L-D2AAK2-Fab3089 complex.**

**a,** Flowchart of Cryo-EM data processing of D_2_R-T4L-D2AAK2-Fab3089 complex.

**b,** Representative cryo-EM micrograph (scale bar, 90 nm).

**c,** Local resolution of final cryo-EM map.

**d,** Angular distribution of all particles in final refinement.

**e,** Golden Standard Fourier Shell Correlation (GSFCS) curve for the final refinement.

**f,** Cryo-EM map and model of the D_2_R-T4L-D2AAK2-Fab3089 complex. Cryo-EM density map and model are shown for all seven transmembrane α-helices, helix 8 and D2AAK2.

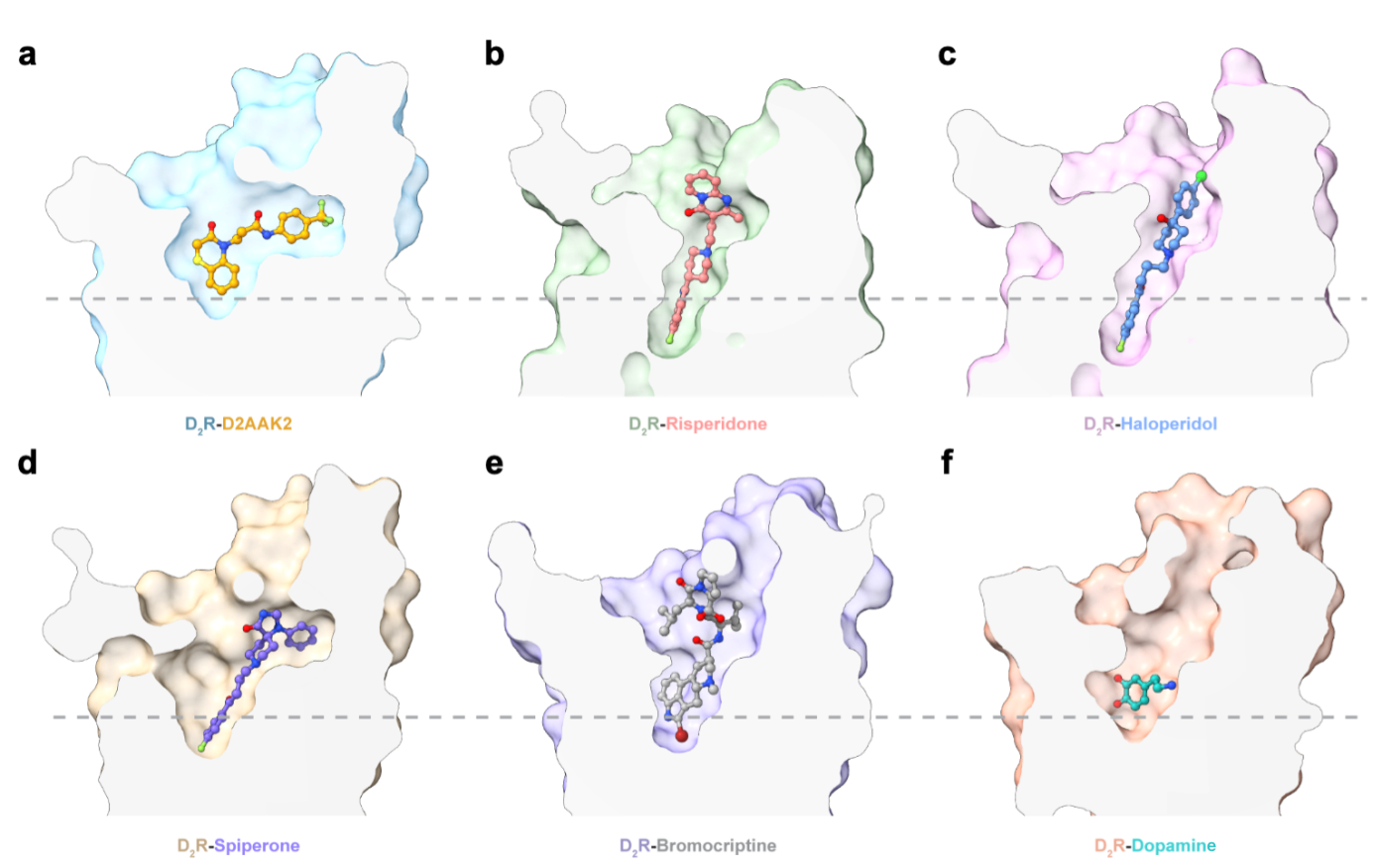

**Fig. S4. Vertical cross sections of the ligand-binding pockets of D_2_R bound with different ligands.** These ligands include antagonists: **a,** D2AAK2; **b,** Risperidone; **c,** Haloperidol; **d,** Spiperone; and agonists: **e,** Bromocriptine; **f,** Dopamine.

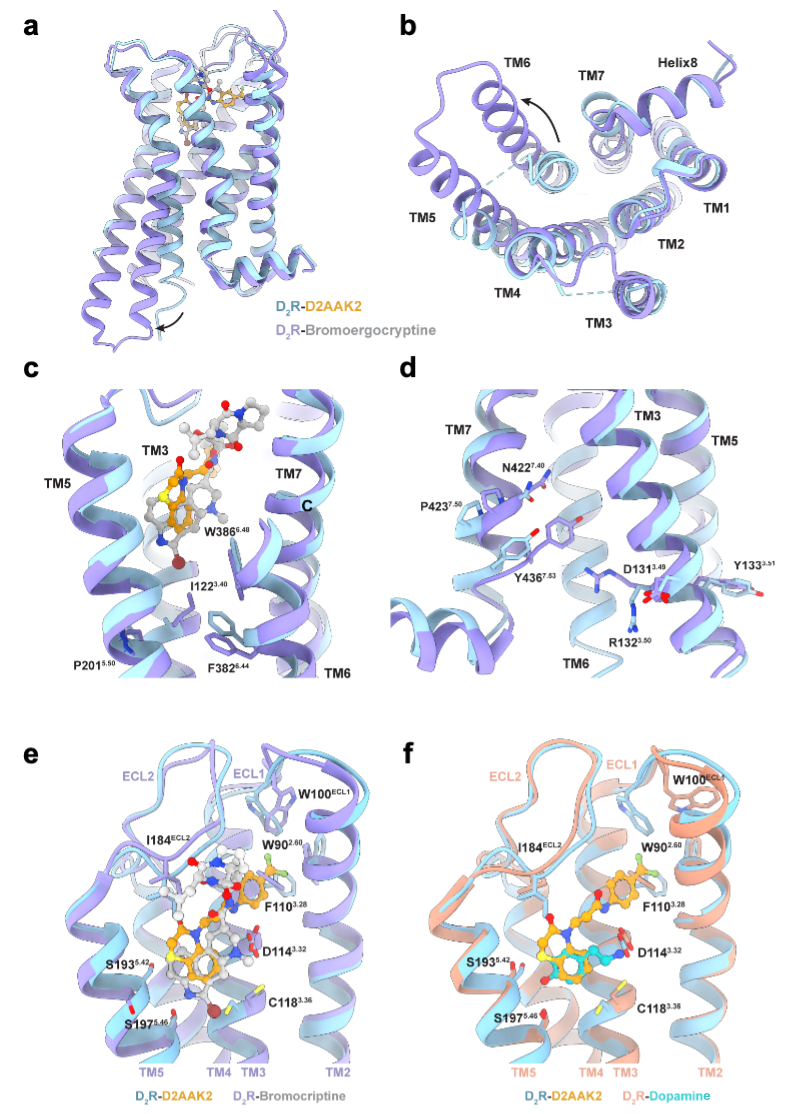

**Fig. S5. Comparison of D2AAK2-bound D_2_R structure with agonist-bound D_2_R structures.**

**a-d,** Structural comparison of the seven transmembrane helices and the activation motifs in the active D_2_R with bromocriptine and the inactive D_2_R with D2AAK2.

**e-f,** Comparison of D_2_R bound with D2AAK2, bromocriptine and dopamine.

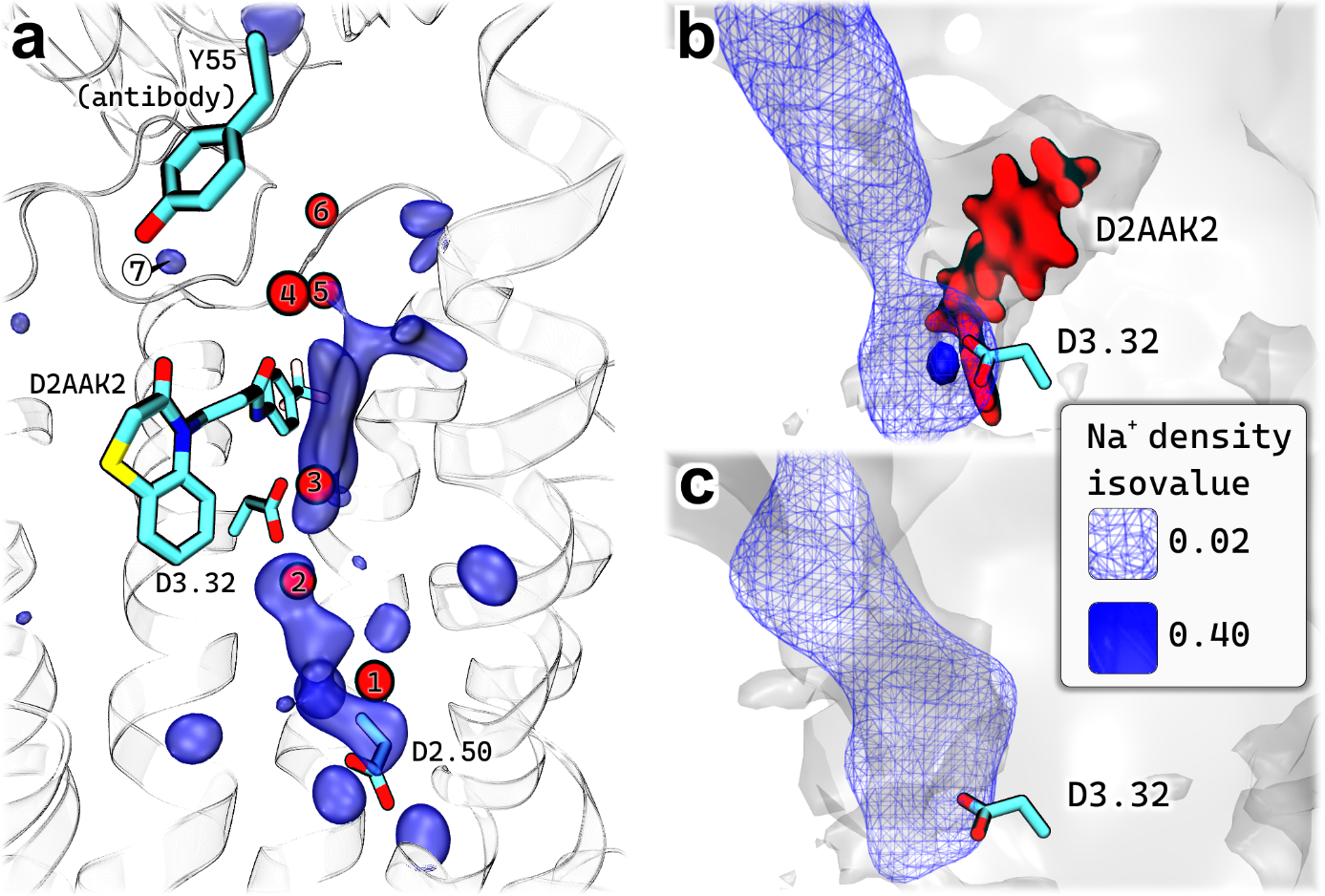

**Fig. S6. Water and sodium occupancy analysis.**

**a,** Water occupancy analysis. Red spheres represent water molecules found in the Cryo-EM structure. Blue transparent surface depicts the water occupancy isosurface.

**b-c,** a comparison of sodium ion density distributions within a receptor cavity in the presence and absence of D2AAK2 in the binding pocket. **b,** the receptor with the bound ligand. **c,** the cavity in the absence of the ligand. A noticeable difference in density isovalues is evident near the D^3.32^ residue, suggesting a potential influence of D2AAK2 binding on the stability of the aspartate-sodium interactions.

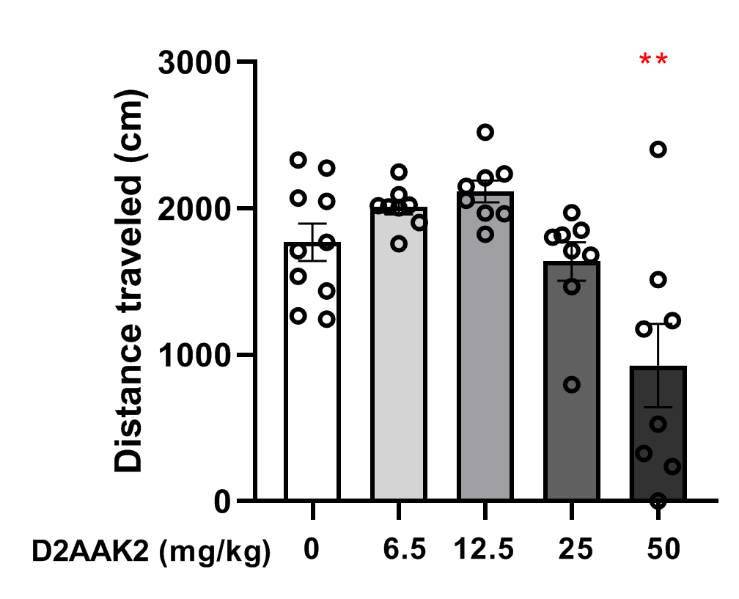

**Fig. S7. Effect of D2AAK2 (6.5, 12.5, 25, and 50 mg/kg, i.p.) on male mouse spontaneous locomotor activity.**

Each dose and vehicle were injected (i.p) 60 min before testing. The data are presented as the mean ± SEM (n = 8-10 animals per group). The results indicated statistically significant difference in distance traveled by animal, the group treated with 50 mg/kg D2AAK2 (i.p.) significantly decreased the time traveled by animals when compared vs control group: **p < 0.01 (One-way ANOVA: F (4, 37) = 8.752, p < 0.0001 followed by Bonferroni’s post hoc test).

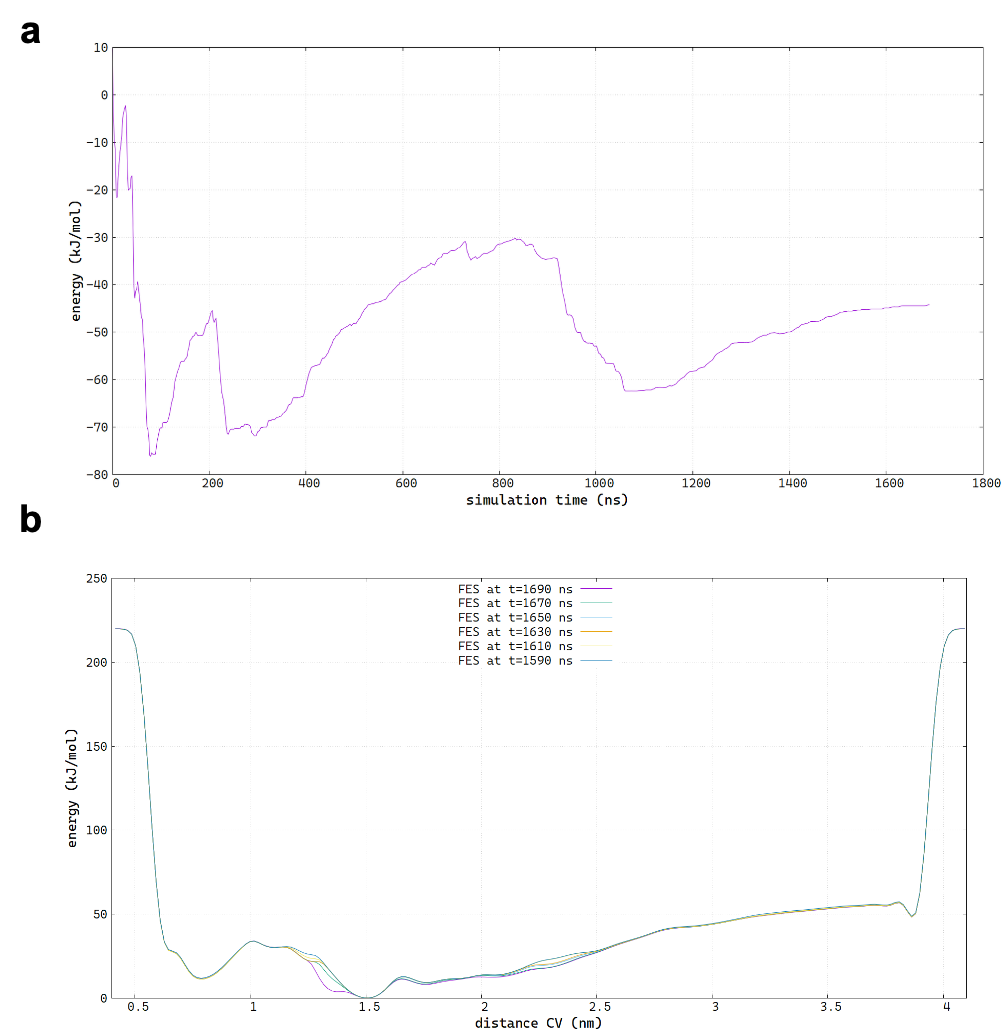

**Fig. S8. Convergence Analysis of Ligand Binding Simulations.**

**a,** Convergence of the FM simulation shown on the plot of ligand binding free energy over simulation time.

**b,** Convergence of the metadynamics simulations shown with snapshots of FES taken every 20 ns for the last 100 ns of the simulation.

**Table S1. Modulation of quinpirole responses by D2AAK2 in cAMP functional data at D_2_R fits globally to the modified Gaddum/Schild EC_50_-shift model of competitive antagonism.** Modified Gaddum/Schild EC_50_-shift model global fitting parameters. Estimated parameter values represent the mean and 95% CI of three independent experiments performed in duplicate or triplicate.

| **Parameter** | **Value** | **95% CI** |
| --- | --- | --- |
| **Log EC_50_**; **p*A*_2_**; **Schild Slope**; **Bottom**; **Top**; **Hill Slope**; **EC_50_**; ***K*_B_** | -8.032; 7.942; 0.8669; -2.406; 82.97; 0.7527; 9.283e-009; 1.143e-008 | -8.198 to -7.868; 7.573 to 8.355; 0.7283 to 1.020; -5.009 to 0.05523; 80.61 to 85.43; 0.6372 to 0.8982; 6.342e-009 to 1.357e-008; 4.413e-009 to 2.674e-008 |

**Table S2. Results for D2AAK2 analogs 1a-d (10 µM) in competition radioligand binding assays at D_2_R.** Data (mean ± SEM of the indicated number of experiments performed in duplicate) are expressed as % of inhibition of [^3^H]-Spiperone specific binding (%inh.). %inh. for haloperidol (1 µM; included as control in these assays) was 99 ± 2% (*n = 5*).

| **Compound** | ***h*D_2_ (%inh. at 10 µM)** |
| --- | --- |
| **1a** | 7 ± 2% *(n = 3)* |
| **1b** | 10 ± 0% *(n = 2)* |
| **1c** | -1 ± 4% *(n = 3)* |
| **1d** | 14 ± 8% *(n = 2)* |

**Table S3.** Statistical analysis showing D2AAK2 effect (6.5, 12.5, and 25 mg/kg) on amphetamine-induced animal hyperactivity.

| **Two-way ANOVA analysis** | | | | |
| --- | --- | --- | --- | --- |
| **D2AAK2 effect on amphetamine-induced animal hyperactivity** | | | | |
| **Dose of D2AAK2** | **Pretreatment effect with D2AAK2** | **Amphetamine treatment effect** | **ANOVA Interaction** | **Post hoc**  **(**Tukey's multiple comparisons test) |
| 6.5,  12.5, and  25 mg/kg | F (3, 62) = 3.604,  p = 0.0182 | F (1, 62) = 130.8,  p < 0.0001 | F (3, 62) = 3.616, p = 0.0179 | VEH vs amphetamine  p < 0.0001  6.5 mg/kg D2AAK2 + amphetamine vs  amphetamine  ns  12.5 mg/kg D2AAK2 + amphetamine vs  amphetamine  p = 0.0489  25 mg/kg D2AAK2 + amphetamine vs  amphetamine  p = 0.0011 |

**Table S4: Detailed experimental conditions employed in competition radioligand binding assays.**

|  | *h*D_2_ | *h*D_1_ | *h*D_3_ | *h*5-HT_1A_ | *h*5-HT_2A_ | *h*5-HT_7_ | *h*H_1_ | *h*M_1_ |
| --- | --- | --- | --- | --- | --- | --- | --- | --- |
| Protein/well | 10 µg | 4 µg | 1 µg | 20 µg | 120 µg | 2 µg | 30 µg | 10 µg |
| Assay buffer | 50 mM Tris-HCl, 120 mM NaCl, 5 mM KCl, 5 mM MgCl_2_, 1 mM EDTA (pH = 7.4) | 50 mM Tris-HCl, 5 mM MgCl_2_ (pH = 7.4) | 50 mM Tris-HCl, 5 mM MgCl_2_ (pH = 7.4) | 50 mM Tris-HCl, 5 mM MgSO_4_ (pH = 7.4) | 50 mM Tris-HCl  (pH = 7.5) | 50 mM Tris-HCl, 4 mM MgCl_2_, 1 mM ascorbic acid, 0.1 mM pargyline (pH = 7.4) | 41.4 mM Na_2_HPO_4_, 8.6 mM KH_2_PO_4_ (pH = 7.5) | 50 mM HEPES, 5 mM MgCl_2_, 1 mM CaCl_2_, 0.2% BSA (pH = 7.4) |
| Radioligand | 0.2 nM [^3^H]- Spiperone^a^ | 0.7 nM [^3^H]- SCH-23390^b^ | 1 nM [^3^H]- Spiperone^a^ | 2 nM [^3^H]-8-OH-DPAT^c^ | 1 nM [^3^H]-Ketanserin^d^ | 2 nM [^3^H]-SB269970^e^ | 2 nM [^3^H]-Pyrilamine^f^ | 2 nM [^3^H]-Pirenzepine^g^ |
| Nonspecific binding | 10 µM Sulpiride | 1 µM Butaclamol | 1 µM Haloperidol | 10 µM Serotonin | 1 µM Methysergide | 25 µM Clozapine | 10 µM Triprolidine | 200 µM Pirenzepine |
| Incubation | 25ºC/120 min | 27ºC/60 min | 25ºC/60 min | 37ºC/120 min | 37ºC/30 min | 37ºC/60 min | 27ºC/60 min | 25ºC/90 min |
| Filter plate | GF/C^h^ | GF/C^h^ | GF/C^h^ | GF/C^h^ | GF/B^i^ | GF/C^h^ | GF/B^i^ | GF/C^h^ |
| Filter plate pre-treatment | presoaked with 0.5% PEI^j^ 1 h and washed with assay buffer | presoaked with 0.5% PEI^j^ 1 h and washed with assay buffer | presoaked with 0.5% PEI^j^ 1 h and washed with assay buffer | presoaked with 0.5% PEI^j^ 1 h and washed with assay buffer | presoaked with 0.5% PEI^j^ 1 h and washed with assay buffer | presoaked with assay buffer | presoaked with 0.1% Tween20 1 h and washed with assay buffer | presoaked with 0.5% PEI^j^ 1 h and washed with assay buffer |
| Wash buffer | 50 mM Tris-HCl, 0.9% NaCl (pH = 7.4) | 50 mM Tris-HCl (pH = 7.4) | 50 mM Tris-HCl (pH = 7.4) | 50 mM Tris-HCl  (pH = 7.4) | 50 mM Tris-HCl  (pH = 6.6) | 50 mM Tris-HCl, 4 mM MgCl_2_, 1 mM ascorbic acid, 0.1 mM pargyline (pH = 7.4) | 41.4 mM Na_2_HPO_4_, 8.6 mM KH_2_PO_4_ (pH = 7.5) | 50 mM HEPES, 500 mM NaCl, 0.1% BSA (pH = 7.4) |
| Washing steps | 4 x 250 µl wash buffer | 4 x 250 µl wash buffer | 4 x 250 µl wash buffer | 4 x 250 µl wash buffer | 6 x 250 µl wash buffer | 4 x 250 µl wash buffer | 6 x 250 µl wash buffer | 4 x 250 µl wash buffer |

^a^ 76.1 Ci/mmol, 1 mCi/ml, PerkinElmer NET1187250UC. ^b^ 73.1 Ci/mmol, 1 mCi/ml, PerkinElmer NET930250UC. ^c^ 135.2 Ci/mmol, 1 mCi/ml, PerkinElmer NET929250UC. ^d^ 50.3 Ci/mmol, 1 mCi/ml, PerkinElmer NET791250UC. ^e^ 45.8 Ci/mmol, 0.25 mCi/ml, PerkinElmer NET1198U250UC. ^f^ 20 Ci/mmol, 1 mCi/ml, PerkinElmer NET594250UC. ^g^ 85.0 Ci/mmol, 1 mCi/ml, PerkinElmer NET780250UC. ^h^ MultiScreen_HTS_ FB Filter Plate, PerkinElmer MSFBN6. ^i^ MultiScreen_HTS_ FB Filter Plate, PerkinElmer MSFCN6. ^j^ PEI, Polyethyleneimine.

**Table S5.** Cryo-EM data collection, reﬁnement, and validation statistics

|  | D2R-D2AAK2-Fab3089  EMD-67996;  PDB: 21TZ |
| --- | --- |
| **Data collection and processing** | |
| Magnification | 81,000 |
| Voltage (kV) | 300 |
| Electron exposure (e^–^/Å^2^) | 50 |
| Defocus range (μm) | -1.3~-1.6 |
| Pixel size (Å) | 1.0825 |
| Symmetry imposed | C1 |
| Initial particle images (no.) | 1,812,899 |
| Final particle images (no.) | 253,082 |
| Map resolution (Å) | 2.99 |
| FSC threshold | 0.143 |
| Map resolution range (Å) | 2.53-8.73 |
| **Model** | |
| Initial model used | 7DFP |
| (PDB code) |  |
| Model resolution (Å) | 3.12 |
| FSC threshold | 0.5 |
| Model resolution range (Å) | 2.9-3.1 |
| Map sharpening B factor (Å^2^) | -131.5 |
| **Model composition** | |
| Non-hydrogen atoms | 3821 |
| Protein residues | 486 |
| Ligands | 1 |
| *B* factors (Å^2^) |  |
| Protein | 37.87 |
| Ligand | 31.83 |
| **RMSDs** | |
| Bond lengths (Å) | 0.003 |
| Bond angles (°) | 0.527 |
| **Validation** | |
| MolProbity score | 1.5 |
| Clashscore | 4.47 |
| Poor rotamers (%) | 0 |
| **Ramachandran plot** | |
| Favored (%) | 96.01 |
| Allowed (%) | 3.99 |
| Disallowed (%) | 0 |

**References**

1.	Kaczor, A. A. *et al.* N-(3-{4-[3-(trifluoromethyl)phenyl]piperazin-1-yl}propyl)-1H-indazole-3-carboxamide (D2AAK3) as a potential antipsychotic: In vitro, in silico and in vivo evaluation of a multi-target ligand. *Neurochem. Int.* **146**, 105016 (2021).

2.	Leach, K., Sexton, P. M. & Christopoulos, A. Quantification of allosteric interactions at G protein-coupled receptors using radioligand binding assays. *Curr. Protoc. Pharmacol..* **Chapter 1**, Unit 1.22 (2011).

3.	Żuk, J. *et al.* Allosteric modulation of dopamine D2L receptor in complex with Gi1 and Gi2 proteins: the effect of subtle structural and stereochemical ligand modifications. *Pharmacol Rep.* **74**, 406–424 (2022).

4.	Kaczor, A. A. *et al.* Structure-Based Virtual Screening for Dopamine D2 Receptor Ligands as Potential Antipsychotics. *ChemMedChem* **11**, 718–729 (2016).

5.	Motulsky, H.; Christopoulos, A. Fitting Models to Biological Data Using Linear and Nonlinear Regression; GraphPad Software Inc.: San Diego, 2003.

6.	Kaczor, A. A. *et al.* In vitro, molecular modeling and behavioral studies of 3-{[4-(5-methoxy-1H-indol-3-yl)-1,2,3,6-tetrahydropyridin-1-yl]methyl}-1,2-dihydroquinolin-2-one (D2AAK1) as a potential antipsychotic. *Neurochem. Int.* **96**, 84–99 (2016).
